# Lightness illusions and saliency maps for reflectance estimation are similar across a wide range of deep neural network architectures

**DOI:** 10.64898/2026.11.19.767848

**Authors:** Jaykishan Patel, Alban Flachot, Javier Vazquez-Corral, David H. Brainard, Thomas S. A. Wallis, Richard F. Murray

**Affiliations:** Department of Psychology and Centre for Vision Research, York University; Department of Computer Science, Justus Liebig University Giessen, Germany; Department of Computer Science, Universitat Autònoma de Barcelona; Computer Vision Center, Barcelona; Department of Psychology, University of Pennsylvania; Centre for Cognitive Science and Institute for Psychology, Technical University of Darmstadt, Germany; Center for Mind, Brain, and Behavior (CMBB), Universities of Marburg, Giessen, and Darmstadt, Germany

## Abstract

People perform many visual tasks that depend on having representations that remain correlated with surface reflectance despite large variations in illumination and context. This ability is known as ‘lightness constancy’. Developing image-computable models of lightness constancy has been challenging, but we recently showed that deep neural networks trained for reflectance estimation achieve good performance on some lightness tasks, and furthermore are susceptible to many of the same lightness illusions as human observers. Here we extend this work by testing whether these findings are specific to the network architecture we used in our previous work, or whether they generalize broadly across architectures and training procedures. We evaluated a range of architectures (ResUNet, VGGUNet, MobileUNet, and Dense Prediction Transformer) that we trained to estimate reflectance from rendered images of three-dimensional scenes. We compared networks where the initial, stimulus-encoding phase used pre-trained weights learned for other tasks, and networks where all weights were optimized for reflectance estimation. We found that all architectures and training procedures tested produced similar reflectance estimation performance on held-out images, and were susceptible to several classic lightness illusions. Furthermore, saliency analyses showed that all networks had broadly similar spatial sensitivity across input images. Together, these results suggest that lightness constancy and susceptibility to lightness illusions do not depend strongly on details of network architecture and training, but instead often emerge robustly in networks that have been optimized to infer surface reflectance from luminance.

## Introduction

Human vision begins with two-dimensional retinal images, and from this starting point the visual system creates representations that are correlated with important properties of three-dimensional scenes and objects, such as surface colour and shape. An important example of this process is *lightness perception*, the visual system’s remarkable ability to create representations that are correlated with achromatic surface reflectance (i.e., proportion of light reflected), despite large variations in lighting and context. Inferring surface reflectance from retinal luminance is highly underconstrained, as many combinations of lighting, surface reflectance, and shape can give rise to the same luminance image (Belhumeur, Kriegman, & Yuille, 1999). Nevertheless, the human visual system can usually, at a glance, generate a percept that is reliably correlated with reflectance over a wide range of viewing conditions.

Just as interesting as the successes of human lightness perception are its dramatic failures. Lightness illusions show that lightness percepts (i.e., perceived reflectance) can differ substantially from true surface reflectance, and these discrepancies may reveal assumptions and heuristics that support visual processing. Illusions have played an important role in the development of theories of lightness perception (Adelson, 1993, 2000; DeValois & DeValois, 1988; Hess & Pretori, 1894/1970; Katz, 1935; Koffka, 1935; White, 1979), and they provide a strong test of quantitative models.

Several approaches have been proposed for understanding lightness perception, including anchoring theories, spatial filtering approaches, and Bayesian formulations (Allred & Brainard, 2013; Brainard et al., 2006; Dakin & Bex, 2003; Land & McCann, 1971). These theories have been successful in explaining several aspects of human perception and performance, often within limited stimulus domains (Adelson, 2000; Gilchrist, 2006; Kingdom, 2008; Murray, 2021). However, a robust, general-purpose, image-computable model of lightness perception has remained elusive.

Recent work in computer vision has shown that deep neural networks (DNNs) can be trained to estimate surface reflectance in realistic, complex scenes, and this raises the question of whether these networks are a starting point for image-computable models of human lightness perception (Boss et al., 2021; Janner, Wu, Kulkarni, Yildirim, & Tenenbaum, 2017; Li, Shafiei, Ramamoorthi, Sunkavalli, & Chandraker, 2020; Sengupta et al., 2019; Yu & Smith, 2018). In previous work, we introduced a convolutional neural network (CNN) that we trained to estimate surface reflectance from achromatic, rendered images of scenes of geometric objects (Patel et al., 2026). This network made good estimates of reflectance within its stimulus domain, and outperformed several previous models of human lightness and brightness perception. Furthermore, the network’s outputs qualitatively reproduced several strong lightness illusions experienced by human observers, again outperforming previous models. These results suggest that, despite some well-known weaknesses as modelling tools (e.g., their large numbers of parameters, and the difficulty of understanding their trained computations; (Wichmann & Geirhos, 2023)), DNNs are nevertheless interesting candidates for image-computable models of human lightness perception.

However, several questions remain about whether DNNs capture the strategies used by the human visual system for lightness perception. Our previous work showed that the ability to estimate reflectance and a susceptibility to illusions emerged from training for reflectance estimation, and that these behaviours largely disappeared when the same architecture was trained for denoising (Patel et al., 2026). Nevertheless, it is unclear to what extent these findings depend on specific architectural choices and training procedures (Gomez-Villa, Martín, Vazquez-Corral, Bertalmío, & Malo, 2020; Saha, Konkle, & Alvarez, 2026). We also showed that, although networks can achieve high levels of lightness constancy, sometimes exceeding human performance, the image regions they rely on can differ substantially from those used by human observers (Flachot et al., 2026). In addition, DNNs vary widely in their architectures and corresponding inductive biases (Ulyanov, Vedaldi, & Lempitsky, 2018), and these differences may affect how different architectures estimate reflectance.

Here we address these questions by evaluating a range of network architectures that we train for reflectance estimation. We include several types of convolutional networks, as well as a transformer-based network (details below). Furthermore, we evaluate two types of training. In one procedure, the initial part of each network (the ‘encoder’) is trained on a non-reflectance-related task (object classification or depth perception), and the rest of the network (the ‘decoder’) is trained to infer reflectance from image features learned by the encoder. In the other procedure, the entire network is trained for reflectance estimation.

We evaluate the trained networks in three ways. First, we assess how accurately they estimate reflectance pixel by pixel, when presented with images that were not used during training. Second, we test whether the networks are susceptible to the lightness illusions we examined in previous work. Third, we use ‘saliency analysis’ to reveal which parts of input images have strong effects on the networks’ outputs (Simonyan, Vedaldi, & Zisserman, 2014; Smilkov, Thorat, Kim, Viégas, & Wattenberg, 2017; Sundararajan, Taly, & Yan, 2017). Saliency analysis is a method developed in the computer vision literature, that evaluates the partial derivative of a network’s output with respect to each pixel of the input. It therefore shows how small, local variations in the input affect the output. It is similar to reverse correlation in physiology and classification images in psychophysics (Ahumada, 2002; Murray, 2011; Ringach & Shapley, 2004).

Together, these experiments aim to determine whether diverse architectures trained for reflectance estimation achieve similar performance, are susceptible to similar lightness illusions, and rely on similar stimulus regions. These are important points for understanding DNNs as models of human lightness perception. They help to answer the question: are specific choices of architecture and training procedure critical for models of lightness, or does human-like perception emerge robustly from a wide range of sufficiently powerful networks, so long as they have been trained to infer reflectance from luminance?

### Related work

As noted above, several theoretical approaches have been proposed for understanding human lightness perception. Thorough reviews are available, including Adelson (2000), Gilchrist (2006), Kingdom (2008, 2011), Brainard and Maloney (2011), Schirillo (2013), and Murray (2021). This work has unquestionably deepened our understanding of lightness perception, and of human vision more generally, but it has not resulted in broadly successful, image-computable models of lightness.

In contrast, in the computer vision literature, DNNs have been successful at estimating reflectance in increasingly complex and challenging scenes. That work builds on a long tradition of computational approaches to intrinsic image decomposition and reflectance estimation (Barrow, Tenenbaum, Hanson, & Riseman, 1978; Shen, Yeo, & Hua, 2013). Shortly after the surge of interest in DNNs due to their success at image classification (Krizhevsky, Sutskever, & Hinton, 2012), Janner et al. (2017) showed that a DNN could simultaneously estimate reflectance, 3D shape, and lighting from images of achromatic objects. Subsequent work extended these approaches to more realistic scenes and materials. For example, Yu and Smith (2018) developed InverseRenderNet, a framework that estimates shape, reflectance, and illumination from a single image using self-supervised learning. Related work demonstrated successful inverse rendering of complex indoor scenes from a single image (Sengupta et al., 2019), and later methods incorporated physically based rendering constraints and improved generalization across diverse scenes and materials (Boss et al., 2021; Li et al., 2020).

Another line of work has shown that training DNNs on low-level image processing tasks, such as denoising, can produce well-known phenomena in lightness and brightness perception as a byproduct. Gomez-Villa et al. (2020) evaluated a range of convolutional architectures trained for denoising and deblurring, and showed that some architectures and training objectives produced brightness and color illusions known from work on human vision. Shallow networks were more likely than deeper networks to produce these illusions, and networks trained for different combinations of denoising and deblurring showed different patterns of illusions despite having similar architectures. This work highlighted the fact that perceptual effects depend not only on network structure, but also on the training task. Related work showed that generative adversarial networks could be used to synthesize novel stimuli that induce lightness illusions in human observers (Gomez-Villa, Martín, Vazquez-Corral, Bertalmío, & Malo, 2022).

Other studies have also explored the relationship between network training objectives and illusion perception. Corney and Lotto (2007) trained a small network to predict surface reflectance in a synthetic ‘dead leaves’ dataset. Even though it had just four hidden units and operated on two-dimensional stimuli, the network accounted for several brightness illusions and predicted parametric variations in illusion strength that were consistent with human perception. Kubota, Hiyama, and Inami (2021) examined shallow convolutional networks trained for denoising, deblurring, or blurring, and compared their outputs to psychophysical data on the Munker-White illusion. They reported mixed results, with the closest matches to human data arising from networks trained to blur images, which contrasts with other findings that emphasize denoising or deblurring objectives. Mukherjee, Paul, and Ghosh (2024) extended this work by comparing neural networks and classical image processing methods for denoising and deblurring. They found that several networks produced brightness illusions, but the classical algorithms they examined did not, suggesting that learned representations, and not merely the task objective, play a critical role in generating illusions. Similarly, Saha et al. (2026) found that several lightness illusions emerged in a network trained to reconstruct achromatic images from the responses of an edge detector.

These studies have shown that deep learning networks trained for low-level image transformations can exhibit lightness and brightness illusions, sometimes in ways that parallel human perception. In most of this previous work the networks were not trained to estimate reflectance, but instead to perform tasks such as deblurring or denoising.

### Saliency maps and interpretability

Understanding why networks behave the way they do is a complementary challenge to optimizing their outputs. In neuroscience and psychophysics, receptive field mapping is a classic approach to understanding complex systems (Ahumada, 2002; Murray, 2011; Ringach & Shapley, 2004), that aims to characterize how local stimulus perturbations affect neuronal or model responses. In the context of artificial neural networks, gradient-based methods are a natural extension of these ideas.

Simonyan et al. (2014) introduced the ‘saliency map’ as a tool for understanding deep networks. In their work, a saliency map is an image that shows the absolute value of the derivative of a network’s scalar output with respect to each input pixel. It indicates how strongly small changes to each input pixel affect the output. Later work developed many alternative forms of saliency analysis, for example evaluating the effect of removing each input pixel completely (i.e., setting its value to zero) instead of the effect of small perturbations (Ancona, Ceolini, Öztireli, & Gross, 2018).

Early saliency methods were developed for image classification networks, and their goal was to show what features made it more likely for a network to assign an image to a given class. Recent work has extended these methods to tasks such as semantic segmentation and depth estimation, where the outputs are images instead of category labels. In these settings, saliency is computed by defining a task-relevant scalar function of the output image, such as the mean over a region of interest, and finding the derivative of this function with respect to each input pixel (Ancona et al., 2018; Selvaraju et al., 2017; Sundararajan et al., 2017). The resulting saliency map indicates which input pixels most strongly influence the output over the region. This approach provides end-to-end saliency maps (Ancona et al., 2018; Arias-Duart, Parés, Garcia-Gasulla, & Gimenez-Abalos, 2022; Von der Haar, Elvira, & Ochoa, 2023). This is the version of saliency analysis we used here, in order to find which image locations had the largest effect on the mean reflectance output of a network over target regions.

### Goals of the present work

Our work builds on previous modeling efforts in three key ways. First, instead of training networks for restoration tasks such as denoising or deblurring, we train them to estimate surface reflectance. This choice reflects the view that lightness perception is shaped by ecological demands, where the visual system must use ambiguous sensory input to produce representations that are correlated with physical properties of objects and scenes. By directly optimizing for reflectance estimation, we test whether this objective results in networks that are susceptible to the same illusions as human vision.

Second, instead of just one network, we train a range of network architectures, to evaluate whether phenomena such as visual illusions depend on specific architectural choices, or instead emerge robustly from training for reflectance estimation. We include both ‘frozen’ networks, where model weights in the first stages of the network were learned for different tasks such as denoising, and ‘fine-tuned’ networks, where all weights were optimized for reflectance estimation.

Third, we combine evaluation of network behaviour with a gradient-based saliency analysis. We examine not only whether networks are susceptible to lightness illusions, but also which image regions contribute most strongly to these effects. Together, these methods provide a framework for assessing the extent to which deep learning models capture the computational strategies that underlie human lightness perception.

## Methods

### Training images

We used a large set of rendered images for supervised learning. The dataset contained 100,000 training image pairs, 5,000 validation pairs, and 5,000 test pairs, rendered using Blender 2.92 with the EEVEE engine. Each pair consisted of a 256 *×* 256 pixel achromatic luminance image and a corresponding reflectance image (Figure 1).

**Figure 1.**
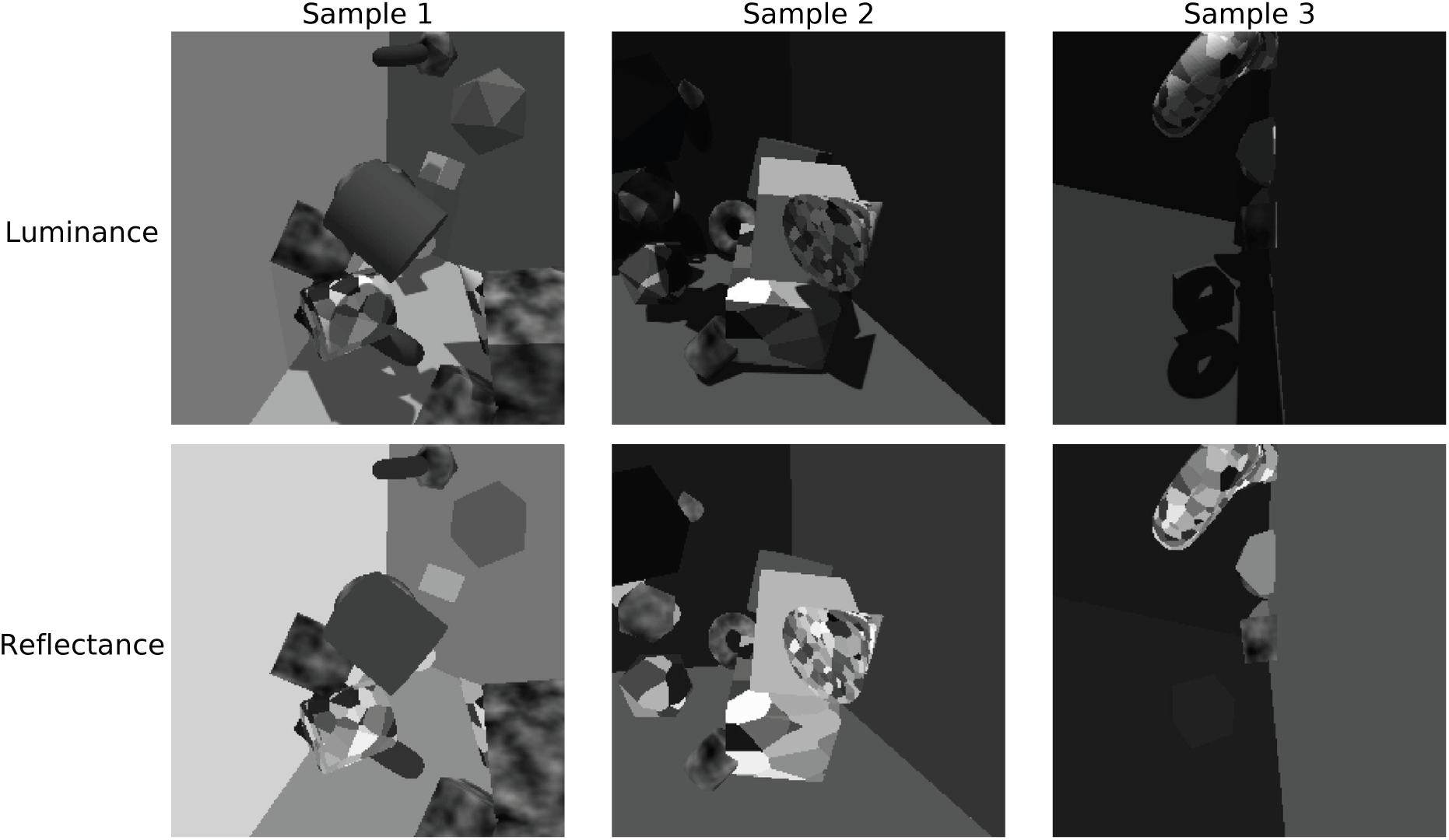
Examples of images used for training and evaluation. Each column shows a rendered three-dimensional scene containing several geometric objects. The top row shows luminance images, and the bottom row shows the corresponding reflectance maps.

This is the same dataset we used in previous work (Patel et al., 2026), except that previously the image resolution was 512 *×* 512, and here we downsampled all images to 256 *×* 256 as that was the resolution required by the transformer-based network.

Blender 2.92 has two render engines: EEVEE, for real-time rasterization, and Cycles, based on ray-tracing. Here we used EEVEE because in previous work we found that Cycles produces a speckle artifact that provides an unintended cue to local illumination, and that networks can learn to exploit this cue (Flachot et al., 2026).

Each rendered scene depicted up to 20 randomly placed geometric objects. Each object was randomly chosen to be a cube, cylinder, sphere, torus, icosahedron, or cone, with position, size, and orientation randomized. Surfaces were Lambertian and achromatic, and each object was assigned one of three reflectance patterns: (1) a uniform grey reflectance randomly sampled from the range [0.05, 0.80], which approximates the range of reflectances observed in natural scenes (Gilchrist, 2006); (2) a randomly generated Voronoi texture; or (3) a low-frequency noise texture. The background of the scene was provided by two walls and a floor, with randomized reflectances and orientations. Illumination consisted of directional and ambient light, with the direction and intensity of the directional source, as well as the intensity of the ambient source, randomized for each scene. Rendering code, including details of the randomization, is provided as Supporting Information (https://github.com/JaykishanPatel/irnet_supporting).

We used rendered images of geometric objects, rather than real or photorealistic images, in order to examine what aspects of lightness perception can emerge from training on images of relatively simple scenes, that nevertheless contain key features such as shadows, shading, and occlusion. This allowed us to test whether aspects of lightness perception can emerge from these generic features, without more specific features of natural scenes such as common objects or materials.

### Network architectures

We evaluated four DNN architectures: three convolutional neural networks (CNNs: VGGUNet, ResUNet, and MobileUNet) and one transformer-based network, the Dense Prediction Transformer (DPT) (He, Zhang, Ren, & Sun, 2016; Howard et al., 2017; Ranftl, Bochkovskiy, & Koltun, 2021; Simonyan & Zisserman, 2015). These architectures differ in their core design principles, including the use of plain convolutional feature hierarchies (VGGUNet), residual connections (ResUNet), lightweight grouped convolutions and squeeze-and-excitation mechanisms (MobileUNet), and a transformer-based architecture with global self-attention and hierarchical patch embeddings (DPT).

Figure 2 illustrates how we used these four architectures. The CNN-based encoders (VGG, ResNet, and MobileNet) were originally designed for image classification, and so produced labels as outputs, not images. To allow them to generate a pixelwise reflectance estimation image, we used these networks as encoders in a U-Net architecture, and appended a decoder that further processed their convolutional layers to produce a reflectance map as an output image. The DPT was originally designed to map input images to output images, so it already had a convolutional decoder and did not require modification.

**Figure 2.**
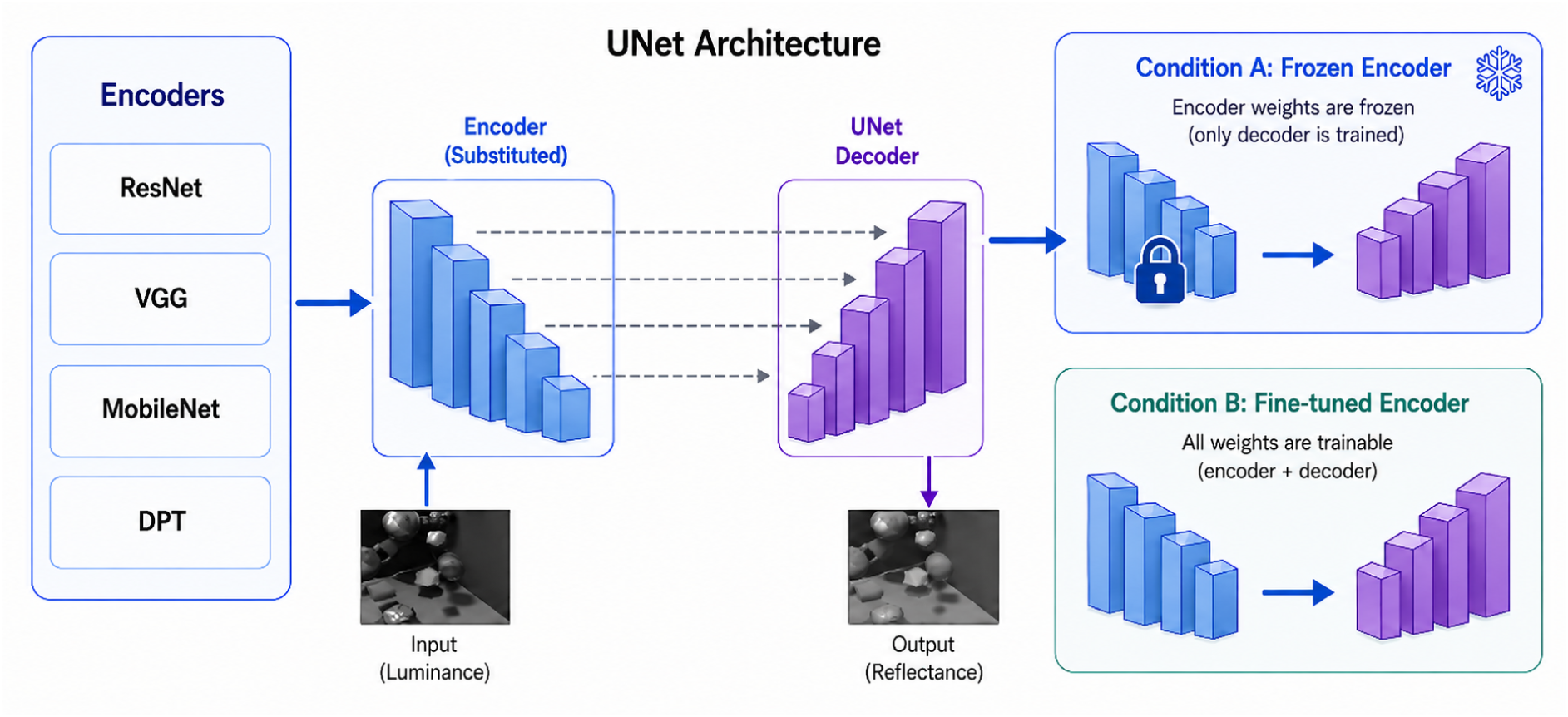
Four encoder architectures (ResNet, VGG, MobileNet, DPT) used an encoder-decoder framework for pixelwise reflectance estimation. The encoder transformed the luminance input into multiscale feature representations, which were passed to a decoder to produce a full-resolution reflectance estimate. Each encoder was evaluated under two training configurations: frozen-encoder, in which encoder weights were frozen and only the decoder was trained, and fine-tuned, in which both encoder and decoder weights were trained. the encoders were fixed at weights learned for ImageNet classification (Deng et al., 2009; He et al., 2016; Howard et al., 2017; Simonyan & Zisserman, 2015). The decoder weights were randomly initialized before training, since these networks were originally trained for image classification and no decoder weights for producing image outputs were available. For the DPT, both the encoder and decoder weights were initialized to values learned for monocular depth estimation (Ranftl et al., 2021). During training, the encoder weights were held fixed and the decoder weights were optimized. This frozen-encoder condition tested the extent to which pretrained visual representations could support reflectance estimation without task-specific modification of the encoder.

This approach allowed us to compare the architectures’ performance at reflectance estimation, despite their different original purposes and internal designs. All networks were configured to accept 256 *×* 256 pixel input images and produce 256 *×* 256 output images.

### Training

Another choice with DNNs is whether to train the network from random initial weights or to incorporate publicly available pretrained weights. Pretrained networks can provide general-purpose visual representations that support a range of tasks, not just the task the network was originally trained on. For example, convolutional encoders trained for image classification have been used successfully in semantic segmentation networks (Iglovikov & Shvets, 2018; Stawiaski, 2019).

For each architecture, we evaluated two training procedures. In the *frozen-encoder* condition, the encoder weights were held fixed at publicly available, pre-trained values, and the decoder weights were optimized during training. For the three CNNs we used, In the *fine-tuned* condition, the weights were initialized in the same way as in the frozen-encoder condition, but during training both encoder and decoder weights were optimized for reflectance estimation.

Networks were trained via supervised learning, using rendered luminance images as inputs and corresponding reflectance maps as target outputs. We optimized mean-squared-error loss using the ADAM optimizer (Kingma, 2015). The batch size was 64 images, and training continued for ten epochs. The learning rate was 10^−3^ for the first 400 training batches and 10^−4^ for the remainder of training. All other optimization parameters were set to the default ADAM values.

Because supervised learning is stochastic, different training runs can yield different learned weights and model behaviours. To characterize this variability, we trained ten model instances for each combination of architecture and training condition. Thus the full design included 4 architectures *×* 2 training conditions (frozen and fine-tuned) *×* 10 independent training runs, yielding 80 trained models.

### Lightness illusions

In addition to the randomly generated test scenes described above, we also evaluated the trained networks on a set of classic lightness illusions (Figure 3). These images were not used during training, and we used them to probe the networks’ responses to images where human observers perceive physically equiluminant regions to have very different reflectances.

**Figure 3.**
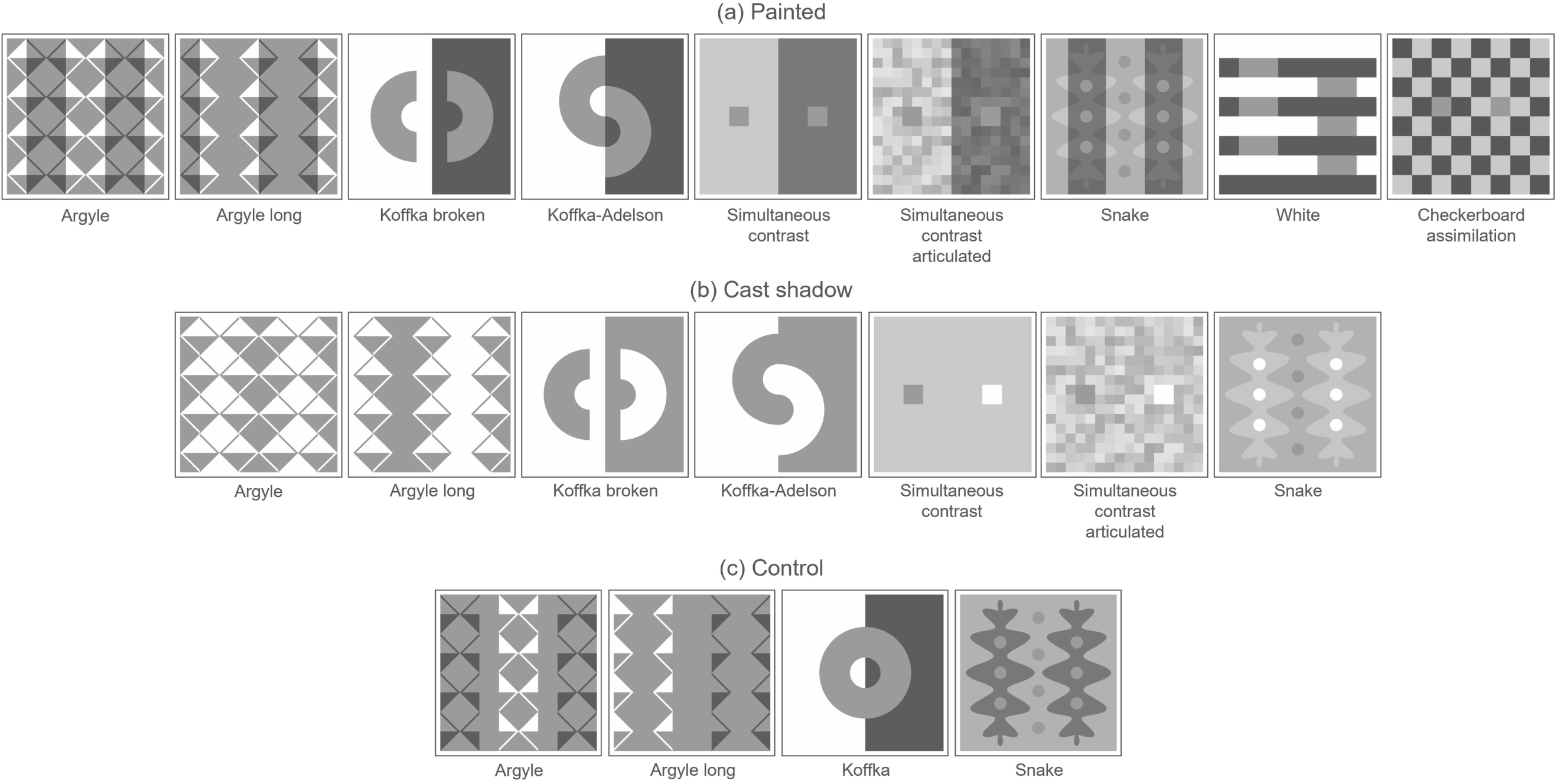
Classic lightness illusions (a) Painted variants, where dark regions are included in the images shown here. (b) Cast-shadow variants, where dark regions were created by cast shadows in a 3D scene (see Figure 5), and are not included in the images shown here. (c) Control variants, where local contrasts are largely preserved but some cues to cast shadows have been eliminated.

We used the argyle illusion and its long-range variant (Adelson, 1993), the Koffka illusion and its control figure (Koffka Broken) (Koffka, 1935), the Koffka-Adelson illusion (Adelson, 2000), a simultaneous contrast illusion (Hess & Pretori, 1894/1970), an articulated simultaneous contrast illusion (Katz, 1935), White’s illusion (White, 1979), the checkerboard assimilation illusion (DeValois & DeValois, 1988), and the snake illusion (Adelson, 2000). In each illusion, two or more regions with the same luminance are perceived by human observers to have different reflectances. We chose regions of interest (ROIs) based on those equiluminant regions (Figure 4). For some illusions (argyle, Koffka, and snake), we included control figures where local luminances surrounding the test patches were largely preserved, but cues to illumination, such as X-junctions, were disrupted. For human observers, the lightness illusion is greatly reduced in these control conditions. This provides a useful test of computational models, as models that rely mostly on local contrast tend to produce similar outputs for the illusion and control figures, unlike human vision. These are the same images we used in previous work (Patel et al., 2026), where we confirmed that human observers perceive the expected illusions in these illusion images, and not in the control images. The images in Figure 3 are very different from the training images in Figure 1.

**Figure 4.**
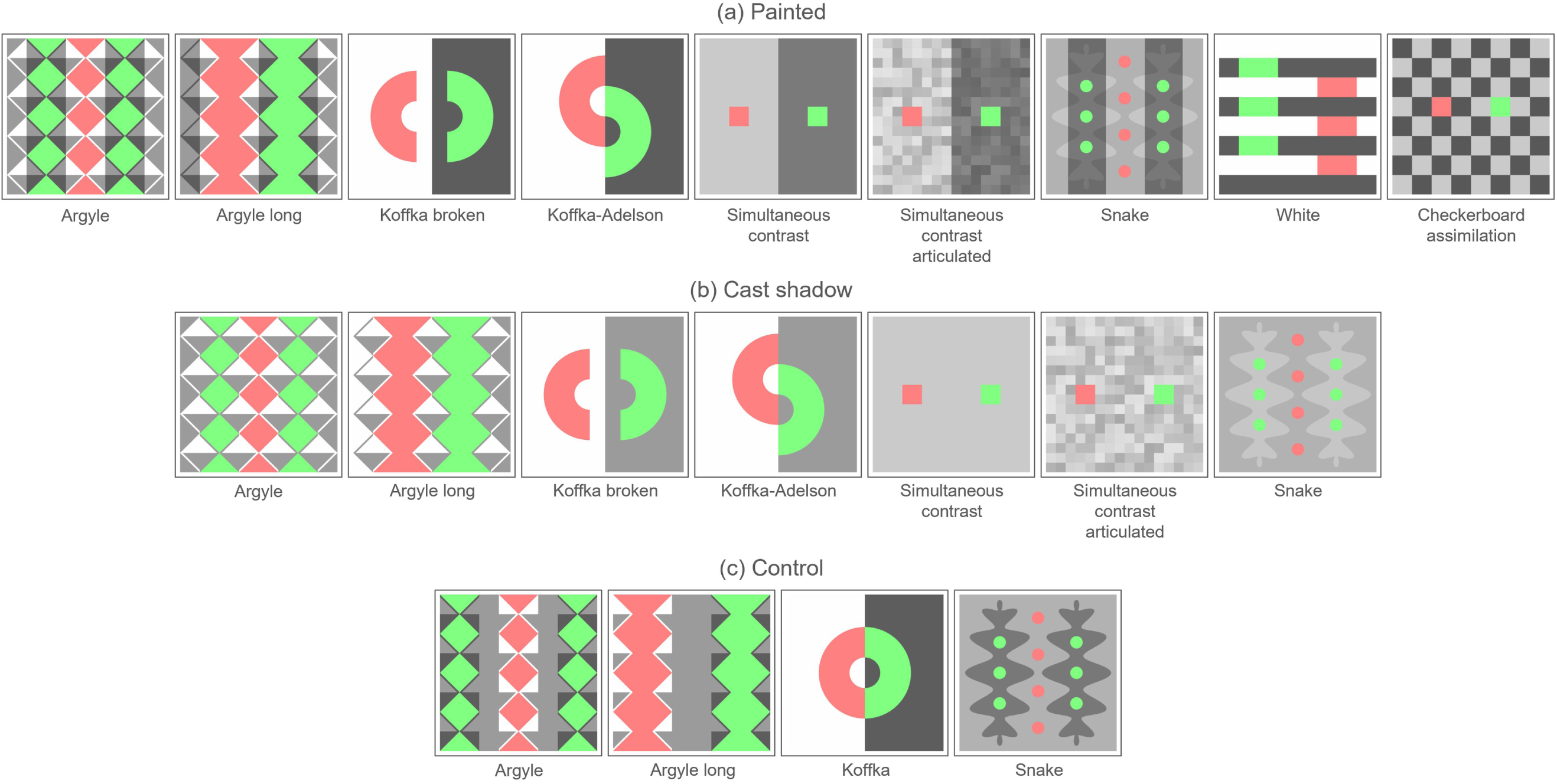
Regions of interest for images shown in Figure 3. Regions indicated in red and green are equiluminant in the corresponding images in Figure 3.

Our main interest was not the ability of networks to generalize far beyond their training set, so we did not test networks directly on the images in Figure 3. Instead, we embedded each of those images in a 3D scene that was more similar to the training set. We mapped each image in Figure 3 onto the faces of a cube in a scene that contained several additional geometric objects (Figure 5). We rendered these 3D images using the same graphics pipeline we used for the training images, as described above.

**Figure 5.**
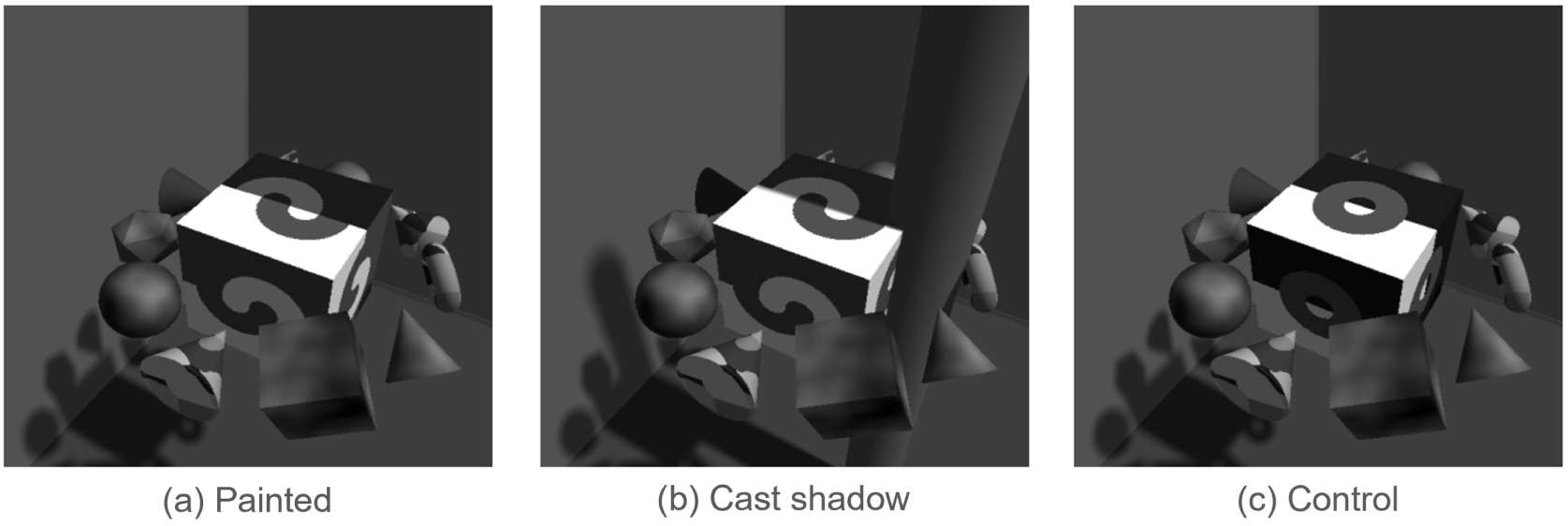
Test images rendered using the Koffka-Adelson illusion and its control variant, as shown in Figure 3.

Shadows are sometimes thought to play an important role in some of these illusions, such as the argyle, Koffka, snake, and simultaneous contrast illusions. Motivated by this idea, we rendered two variants of the images. In the ‘painted’ variant, the illusion was simply rendered on the cube as a surface reflectance pattern (Figures 3a and 5a). In the ‘cast-shadow’ variant, the reflectance image rendered onto the cube lacked certain large, dark reflectance regions, and these dark regions were instead created as shadows cast by one or more upright cylinders in the scene (Figures 3b and 5b). Thus the painted and cast-shadow stimlui were broadly similar, but had some subtle differences, e.g., the cast-shadow stimuli had a small penumbra at the shadow boundary, and included cylinders whose size and position were consistent with the shadows cast on the cube. The control stimuli (Figures 3c and 5c) were created in the same way as the painted stimuli, by simply rendering images onto the sides of cubes as a reflectance pattern.

The lighting configuration and camera parameters were held constant across all illusion stimuli, except for small adjustments in a few cases to ensure that the luminance of the test regions was matched across conditions. Except for a small penumbra at shadow boundaries in the cast-shadow variants, luminance was the same in the ROIs of the painted and cast-shadow versions of each image. All lighting parameters fell within the range used to generate training images.

We quantified the illusion strength for each model by finding the model’s average output over the regions shown as green in Figure 4, minus the average over regions shown as red. In previous work (Patel et al., 2026) we evaluated illusion strength in a different way, using a simulated matching experiment where for each model we found the reflectance that had to be assigned to the match path (red regions) during rendering of the scene, in order for the model to have the same mean output there as at the fixed reference patch (green regions). We tried this approach with the present four architectures, but we found that the illusions were often so strong that no reflectance within the range we used to render the training scenes (0.05 to 0.80) gave a match. This was true for 32% of the attempted matches. Instead of extrapolating the data to find a reflectance match outside the range used for training, we chose to use a different measure of illusion strength, and as explained above, here we report the difference in mean model outputs between the green and red regions in Figure 4.

Another, less common but nevertheless interesting problem that we mention for exploration in future work is that with the present models, outputs at the match patch were sometimes a non-monotonic function of the reflectance of the match patch, so that two different reflectances at the match patch gave equally good matches to the reference patch. This does not seem to be driven by features of the training data, and we speculate that more thorough training would eliminate this problem. In any case, this problem does not affect our present analysis, because instead of using a simulated matching task, we simply report the difference in mean model output over the two target regions that define the illusion (red and green regions in Figure 4).

### Saliency maps

To see which image regions had the strongest effect on model outputs, we computed gradient-based saliency maps (Simonyan et al., 2014). Let *f_θ_* : ℝ*^HW^ →* ℝ*^HW^* represent a trained network with parameters *θ*, that maps an *H × W* pixel luminance image **x** to an equally sized reflectance estimate, ^**r** = *f_θ_*(**x**). For a given input image **x**, each pixel *G_i_* of the saliency map is the absolute value of the derivative of a scalar function *S*(^**r**) of the output with respect to the corresponding pixel *x_i_* of the input:

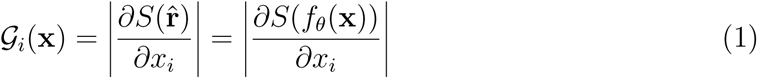

In our first analysis, the scalar objective *S*(^**r**) was the mean reflectance output over all pixels:

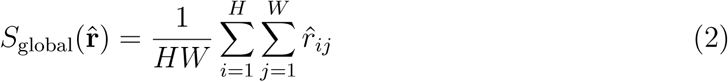

We call this the ‘global’ objective.

In a second analysis, the scalar objective was the mean reflectance output over the regions indicated in green in Figure 4 (which look lighter to human observers in the illusion figures), minus the mean over the red regions (which look darker):

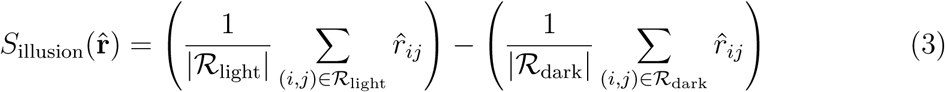

where *|R|* is the number of pixels in a region *R*. We call this the ‘illusion’ objective.

This gradient-based approach provides an end-to-end measure of input sensitivity, and does not examine internal layers. The saliency maps reflect the combined influence of all internal mechanisms, such as encoders, decoders, skip connections, and attention.

## Results

### Reflectance estimation

All models successfully learned to estimate surface reflectance from luminance images. Figure 6 shows samples of luminance images, ground-truth reflectance, and model outputs (predicted reflectance). For all architectures, predicted reflectance closely matched true reflectance on held-out test images, indicating that these results did not depend strongly on our specific architectural choices or training procedure. The networks mostly eliminated shading and shadows, while preserving spatial structure and surface boundaries, and providing broadly accurate pixel-by-pixel estimates of reflectance. This was true with both frozen encoders and fine-tuned encoders.

**Figure 6.**
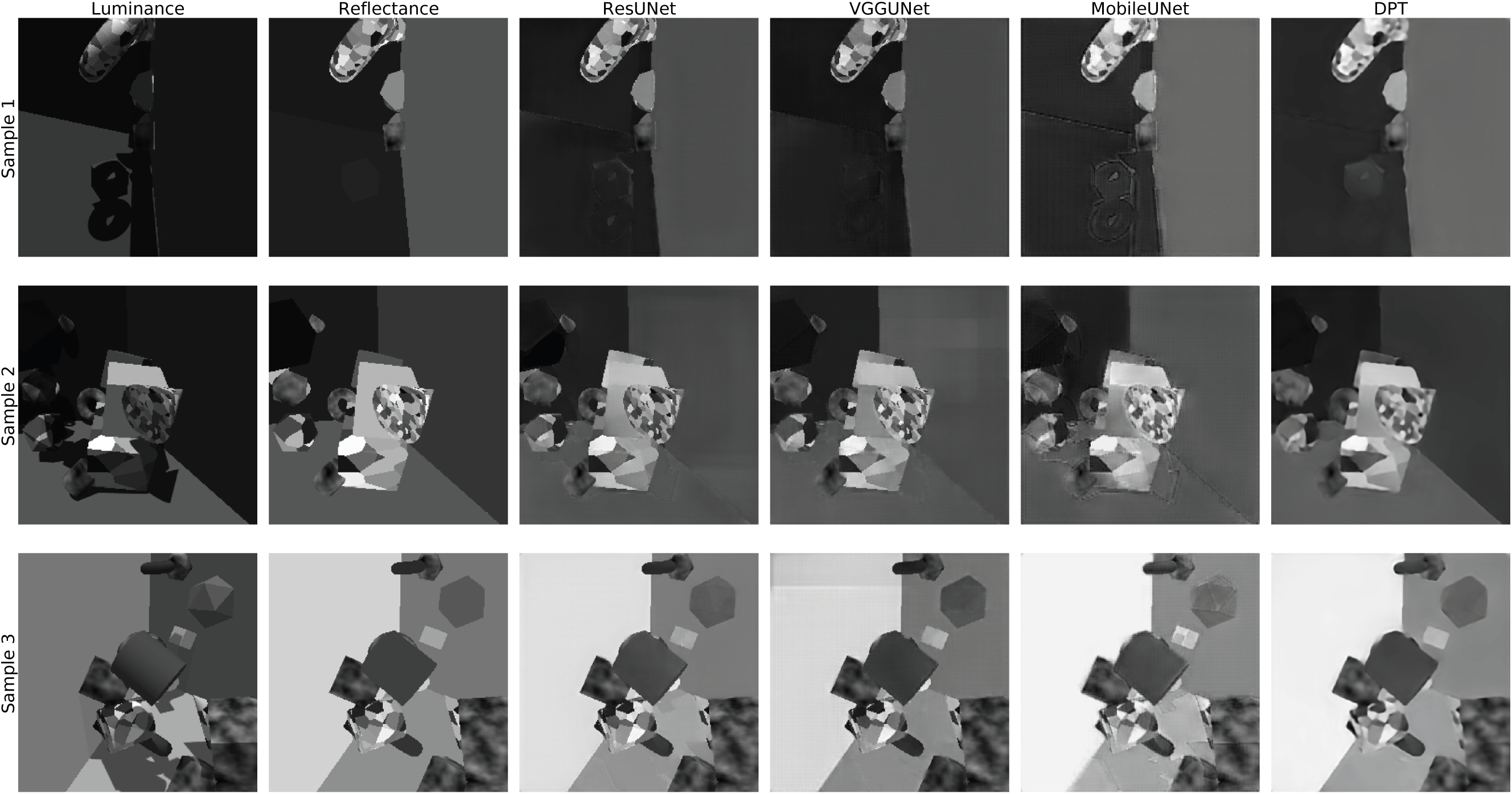
Examples of reflectance estimation results for the fine-tuned models. Each row shows a representative test scene. Columns show the input luminance image, ground-truth reflectance, and the predicted reflectance from each model class. All models recover surface reflectance to some extent while substantially reducing shading and cast shadows. Results with frozen-encoder models were similar.

Figure 7 shows a quantitative assessment of the networks’ performance, plotting predicted versus ground-truth reflectance. Each plot shows results for 10,000 pixels randomly sampled from the 5,000 held-out test images, and averaged across the ten training runs. Across architectures and training conditions, model predictions closely tracked ground-truth reflectance, with strong correlations (*r ≥* 0.92) and low median errors. Fine-tuned models generally had higher correlations and lower errors than their frozen counterparts, indicating that task-specific optimization improves accuracy, though not dramatically.

**Figure 7.**
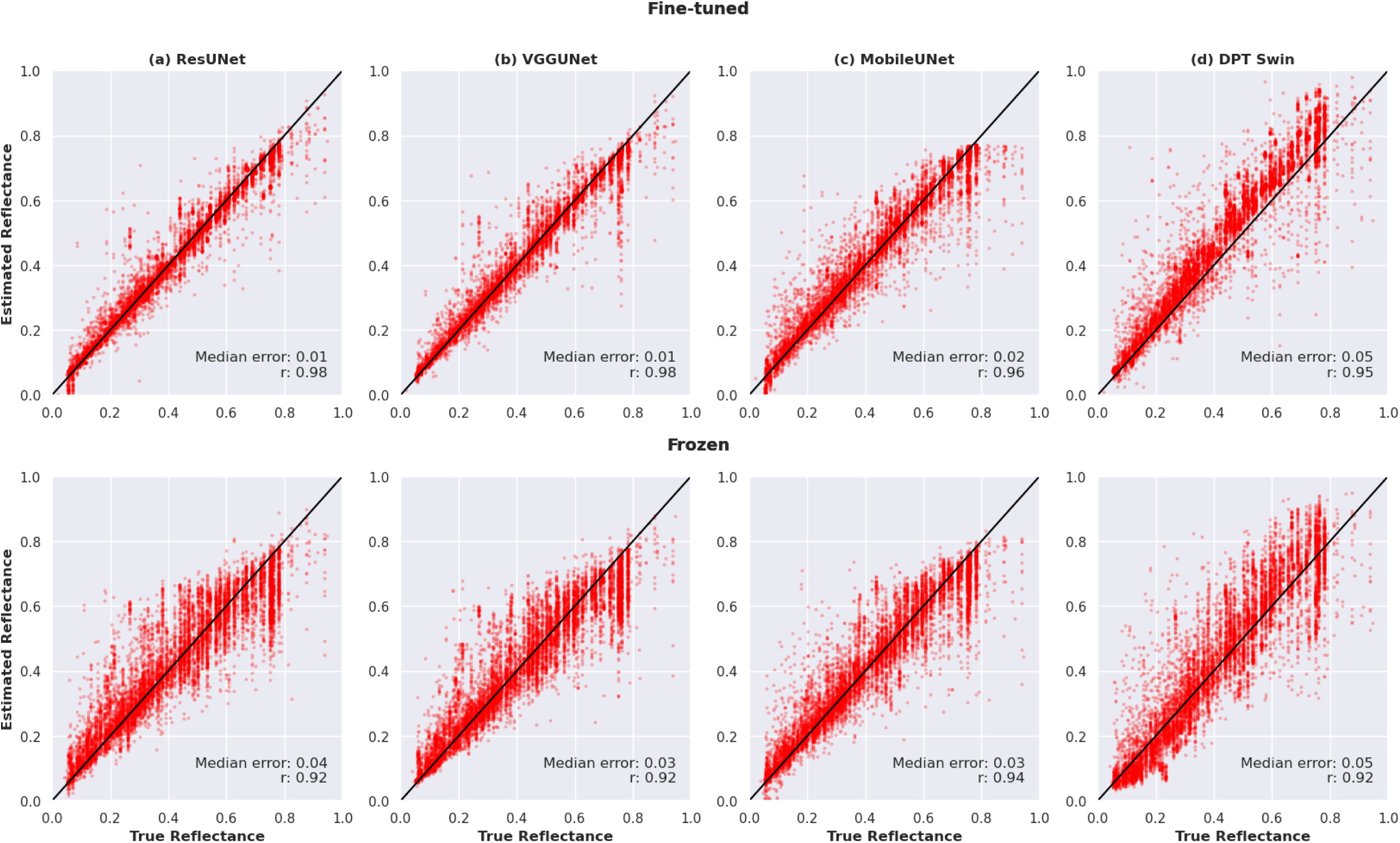
Scatter plots of predicted vs. true reflectance for each architecture and training condition. The top row shows models with fine-tuned encoders, and the bottom row shows models with frozen encoders. Each point represents a randomly sampled pixel from the test set, averaged over the ten training runs. Diagonal black lines represent perfect predictions. Insets report the median absolute reflectance error and Pearson correlation coefficient.

Figure 8 shows another assessment of network performance. For each of the ten independent training runs, we computed the median absolute pixelwise reflectance error across the 5,000 test images, using 100 randomly sampled pixels per image. Figure 8 shows violin plots of the resulting ten median errors for each architecture and training condition. As in the scatter plots discussed above, differences across the highly varied network architectures were relatively modest in absolute terms. With frozen-encoder training, median error ranged from approximately 0.030 to 0.046 reflectance units (on a scale from 0 to 1), whereas with fine-tuned training, it ranged from approximately 0.015 to 0.026. These results suggest that the training data and reflectance estimation objective guided diverse network architectures toward similarly effective reflectance estimation performance. Furthermore, they demonstrate that pre-trained representations for object classification and depth estimation can support reflectance estimation even when the encoders remain frozen, although fine-tuning substantially improves performance. There were, nevertheless, statistically significant differences. A two-way ANOVA showed significant main effects of architecture (F(3, 72) = 9.26, p < effect of training condition was substantially larger than that of architecture. There was also a significant architecture × training-condition interaction (F(3, 72) = 30.01, p < .001, *η*^2^ = 0.556), indicating that the magnitude of improvement with fine-tuning differed across architectures.

**Figure 8.**
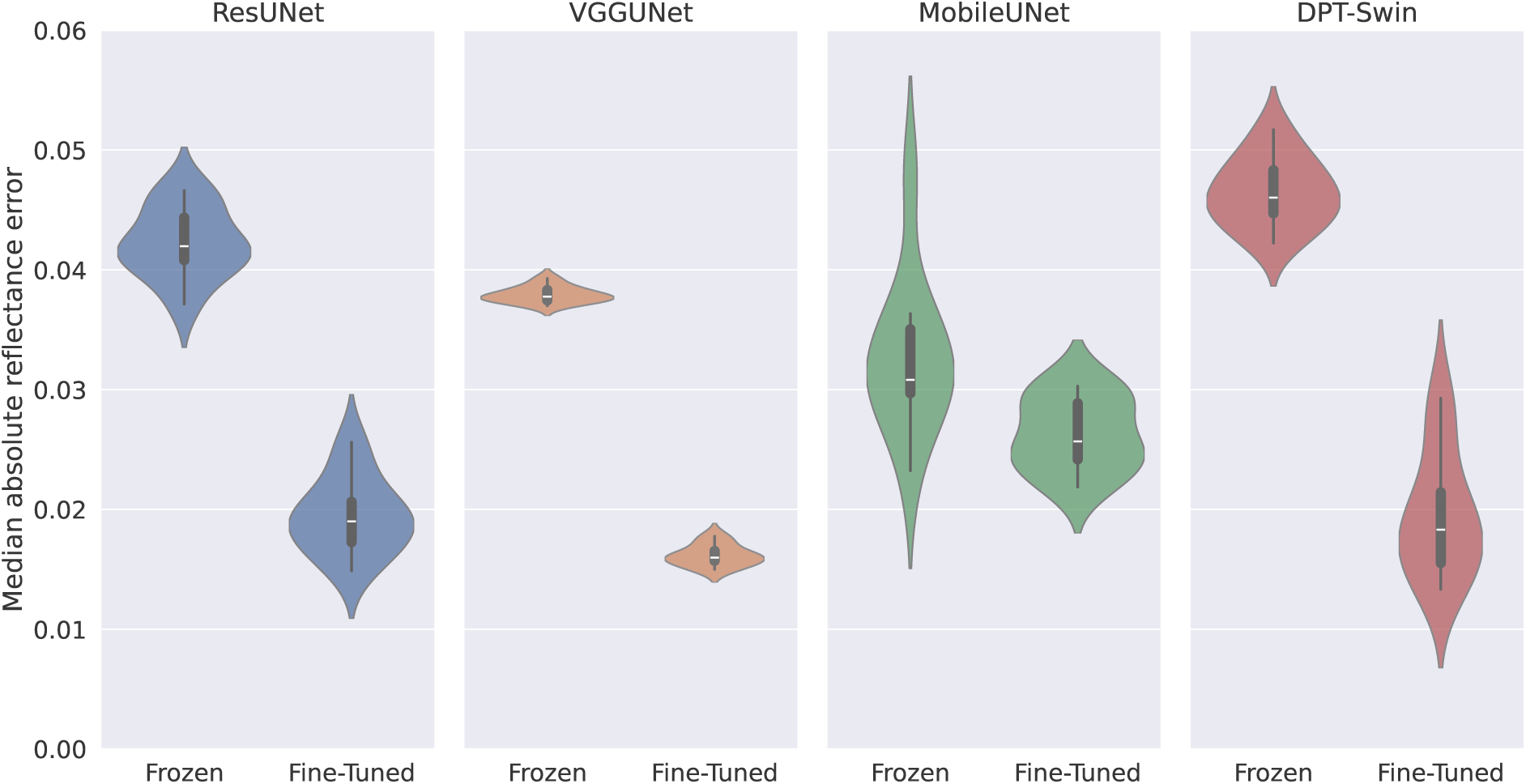
Violin plots of median absolute reflectance error for all models. Fine-tuning reduced error, but good performance was achieved even with frozen encoders.

### Lightness illusions

Figure 9 shows all models’ outputs for one typical illusion (the articulated simultaneous contrast figure). In the input image at the left, the two test squares in the centre of the light and dark halves of the illusion have the same luminance, but to human observers the upper right square (in the dark half) looks lighter. In the model outputs, each model lightens the dark half of the illusion, as if it were partly removing a shadow, and as a result the mean output is higher at the upper right square than at the lower left square, i.e., the model generates the illusion. Discounting of illumination is apparent elsewhere in the outputs too, e.g., the three visible sides of the cube are more similar in the outputs than in the luminance image.

**Figure 9.**
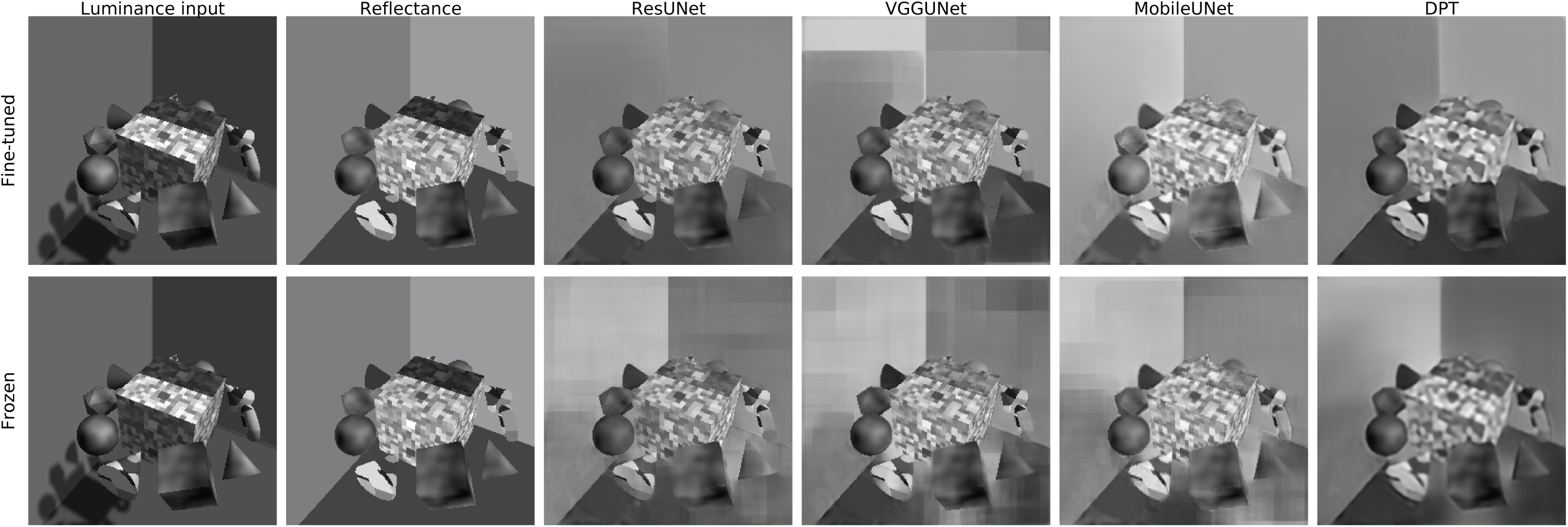
Model outputs for the articulated simultaneous contrast illusion. The two leftmost columns show the luminance input and ground truth reflectance, and the remaining columns show model outputs. The top row shows results for fine-tuned models, and the bottom row shows frozen models.

Figure 10 shows the illusion strength for each model and illusion, averaged over the ten training runs, alongside the corresponding results for human observers. For the models, each bar shows the difference between the model’s average output in the regions coded as green and red in Figure 4. Positive values indicate illusions in the same direction as perceived by human observers. For humans, each bar shows the size of the illusion as measured in a matching task (Patel et al., 2026). For the most part, the models were susceptible to the same illusions as human observers, although the magnitude of the effects varied. This was true for both frozen and fine-tuned models, and for both painted and cast-shadow illusions. Furthermore, illusions that produced relatively strong effects in humans, such as articulated simultaneous contrast, generally also produced strong effects across model architectures, whereas weaker illusions, such as checkerboard, tended to produce smaller model responses. Nevertheless, there were notable differences between human and model responses, both in magnitude and, for some illusions, in direction, as we discuss below.

**Figure 10.**
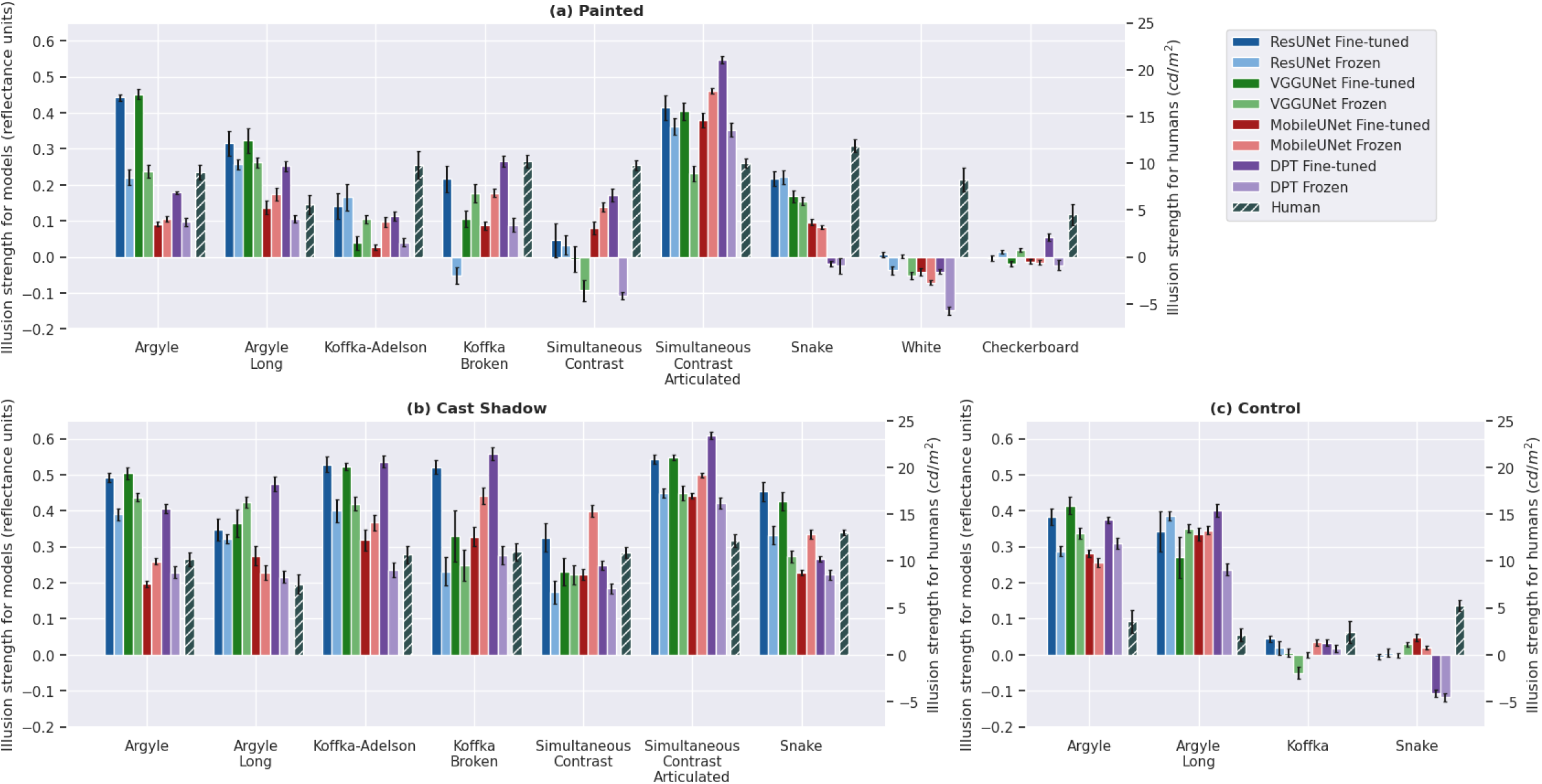
Illusion strength for networks and human observers. (a) Painted, (b) cast-shadow, and (c) control stimuli. For models (solid colors), each bar shows the mean output in the regions indicated by red patches in Figure 4, minus the mean output in regions indicated by green patches; these bars are scaled according to y-axis on the left-hand side of each panel. Each bar shows mean illusion strength across ten training runs, and error bars show standard errors. Positive values indicate illusions in the same direction as perceived by human observers. For humans (striped bars), each bar indicates illusion strength as previously measured in a matching task (Patel et al., 2026); these bars are scaled according to the y-axis on the right-hand side. Each of these bars shows the mean illusion strength across eleven observers, and error bars show standard errors.

In some illusions, target regions surrounded by low luminances appear lighter, and targets surrounded by high luminances appear darker. These are often called ‘contrast’ illusions. The models generally produced strong illusion effects for contrast illusions, including the argyle, long-range argyle, Koffka-Adelson, articulated simultaneous contrast, and snake illusions.

However, for several contrast illusions, the model-generated illusions were weaker in the painted variants than in the cast-shadow variants. Furthermore, for some painted variants (e.g., Koffka Broken, simultaneous contrast, snake), some models even showed near-zero or reversed illusions. In the painted stimuli, the illusions were rendered as reflectance patterns, without cast shadows. The cast-shadow variants had additional cues to illumination, such as penumbras and visible shadow-casting objects.

Accordingly, we might expect a sufficiently powerful network to show no illusions at all for the painted stimuli, where the illusion depends on the observer mistakenly interpreting reflectance edges as shadow boundaries. In that sense, the weaker illusions that we found for painted stimuli represent a success for the networks, not a failure, in terms of their reflectance estimation performance. It does, of course, weaken the networks’ agreement with human judgements.

The control variants of the contrast illusions were designed to disrupt cues to illumination, such as X-junctions, while largely preserving local contrast. The models produced weaker illusions in the Koffka and snake control conditions than in the illusory conditions, which is a success, but they produced strong illusions in the argyle control conditions, which is a failure. Our previous work showed that several classical models fail because they produce equally strong effects in many illusory and control conditions (Patel et al., 2026), so this is a strong test for models of lightness. Here we find that the models we evaluated pass this test, except in the case of the argyle illusion. Previous studies have also found the argyle illusion to be difficult to account for (Kim, Gold, & Murray, 2018; Murray, 2020; Patel et al., 2026).

In a different class of illusions, unlike in contrast illusions, target regions appear lighter when surrounded by high luminances, and darker when surrounded by low luminances. These are called ‘assimilation’ illusions. The models did not account for the two assimilation illusions we examined: for White’s illusion and the checkerboard illusion, most models produced weak or reversed illusions. These results parallel our previous findings, in which assimilation effects have proven more difficult to capture than contrast-based illusions (Patel et al., 2026).

In summary, all architectures, in frozen and fine-tuned variants, exhibited broadly similar patterns of illusions. This indicates that illusion-like effects are robust with respect to choices of architecture or training procedure. They appear to reliably emerge when training networks to infer reflectance from luminance, even in renderings of relatively simple 3D scenes such as those we used here.

### Saliency maps

We start with some qualitative observations about the saliency maps, before reporting a quantitative analysis. Figure 11 shows saliency maps for the global reflectance objective, for the frozen models, for four of the illusions tested, averaged across the ten training runs. The figure shows that pixels that had large effects on the network’s mean output were widely distributed across the image. There were some clear differences between architectures, e.g., ResUNet and VGGUnet had a particularly wide spatial distribution of influential pixels, whereas DPT (the transformer) had more focused clusters. These differences may be related to fundamental architectural differences between the networks, e.g. CNNs are based on local convolutions, whereas transformer networks have a global self-attention mechanism that supports interactions between distant image regions. The saliency maps also show an interesting and unexpected mottling, where even within regions that generally had a strong effect on the output (e.g., the upper left of the ResUNet maps), small groups of pixels that had large effects were separated by thin strips that had much smaller effects. This seems unlikely to be an efficient use of stimulus information for estimating reflectance, and may be worth investigating in later work as a clue to how to improve model performance. These artifacts may be related to quantized features of the network architectures, such as the small kernel size or the sampling stride between layers.

**Figure 11.**
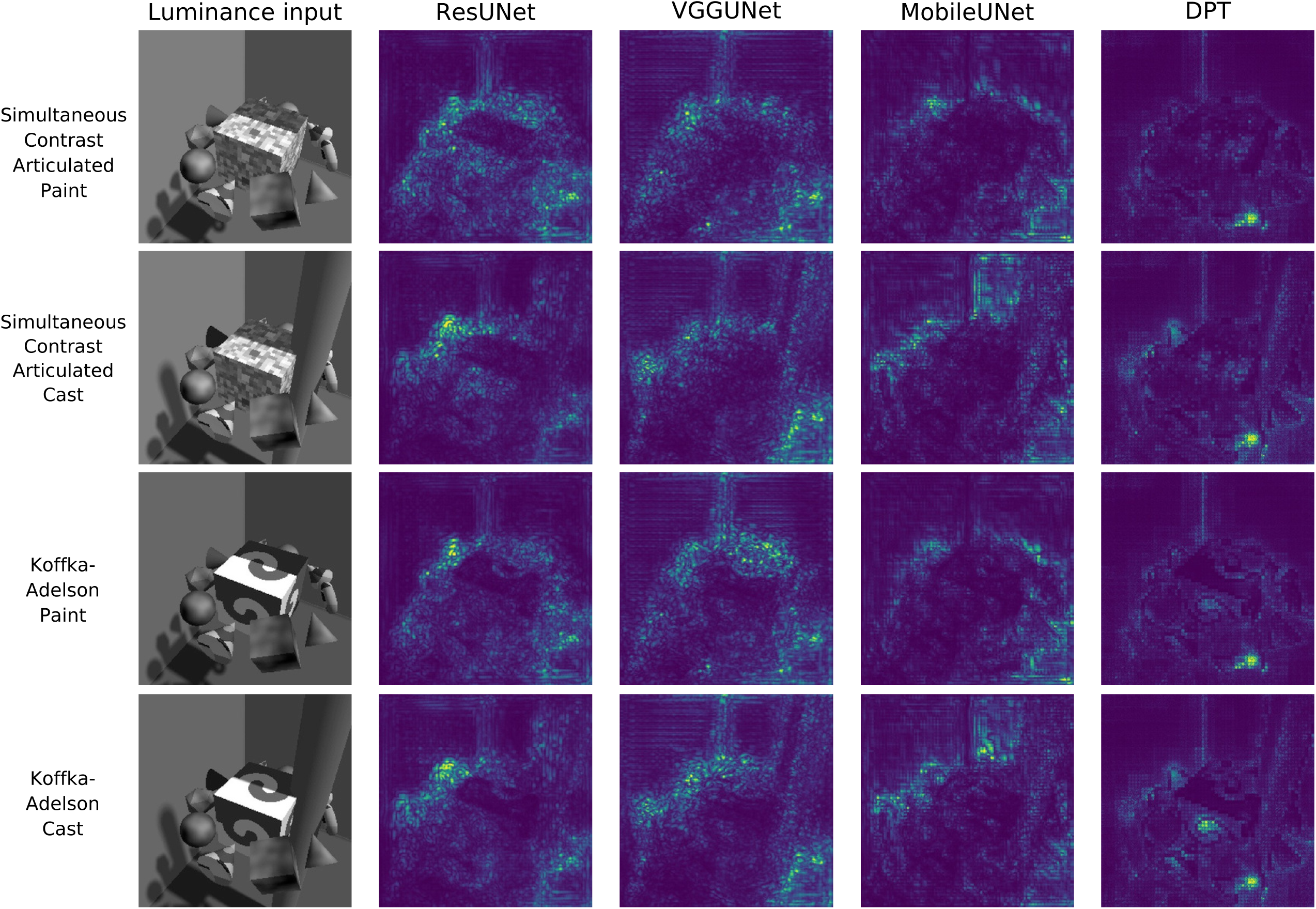
Saliency maps: global objective, frozen models. Columns show results for different architectures.

Figure 12 shows results for the same global objective, for the fine-tuned models. Here influential pixels were still widely distributed across the image, although they were clustered into fewer groups than for the frozen networks. There were again clear differences between architectures. The mottling effect was reduced, but still present, e.g., for VGGUNet, in the cast-shadow conditions, the most influential pixels on the cylindrical columns were organized as thin diagonal stripes.

**Figure 12.**
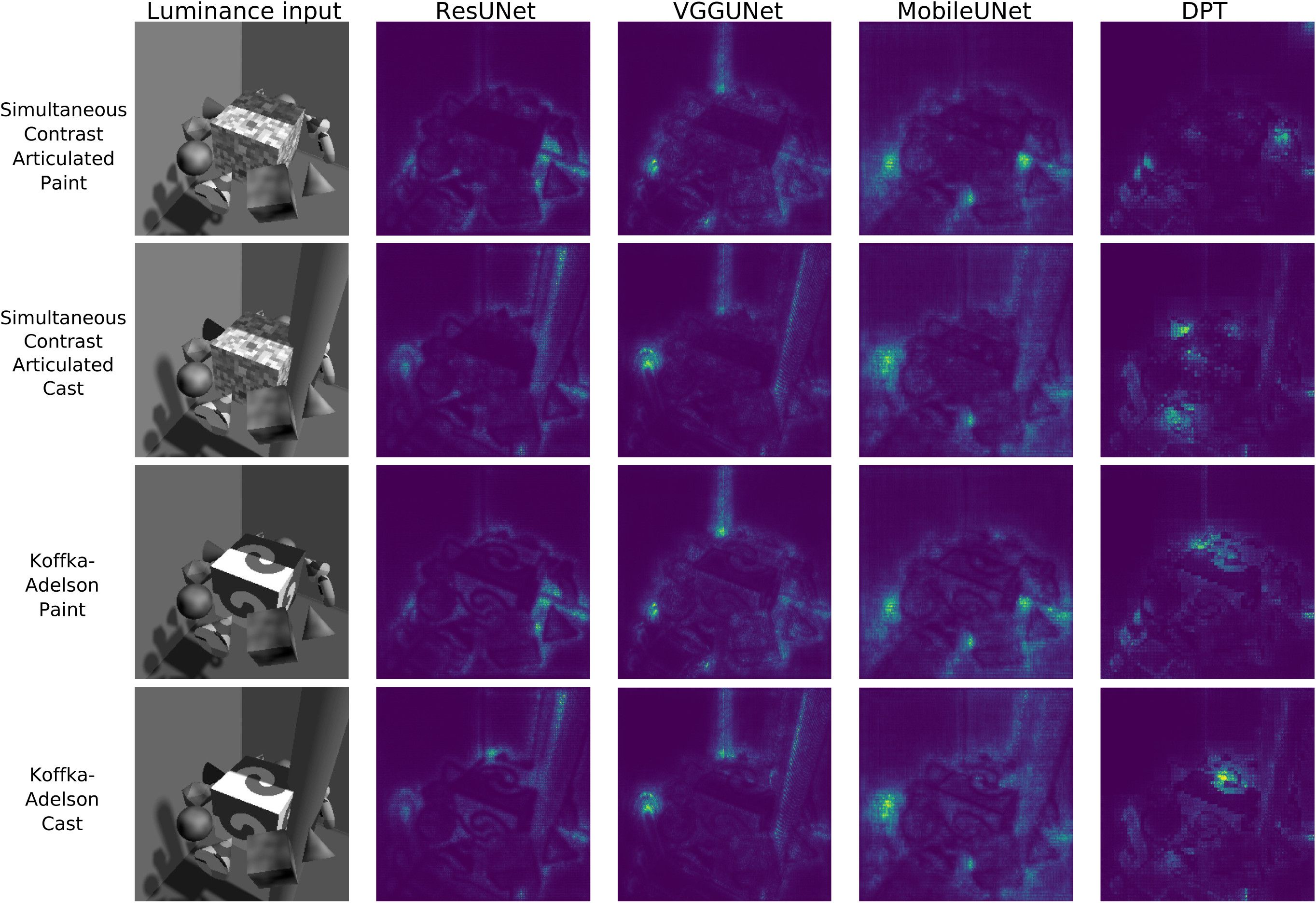
Saliency maps: global objective, fine-tuned models.

Figures 13 and 14 show results for the illusion-based objective, for the frozen and fine-tuned models, respectively. Illusion strength was most sensitive to pixels near the target patches, as we might expect, although nearby pixels outside the target patches often had an influence as well. In illusions that showed light and dark halves of a square, pixels in the darker half tended to have larger effects, e.g., for frozen ResUNet in the simultaneous contrast figures. The differences across architectures and between frozen and fine-tuned networks were less pronounced than for the global objective, although here too there appears to be less mottling for the fine-tuned models.

**Figure 13.**
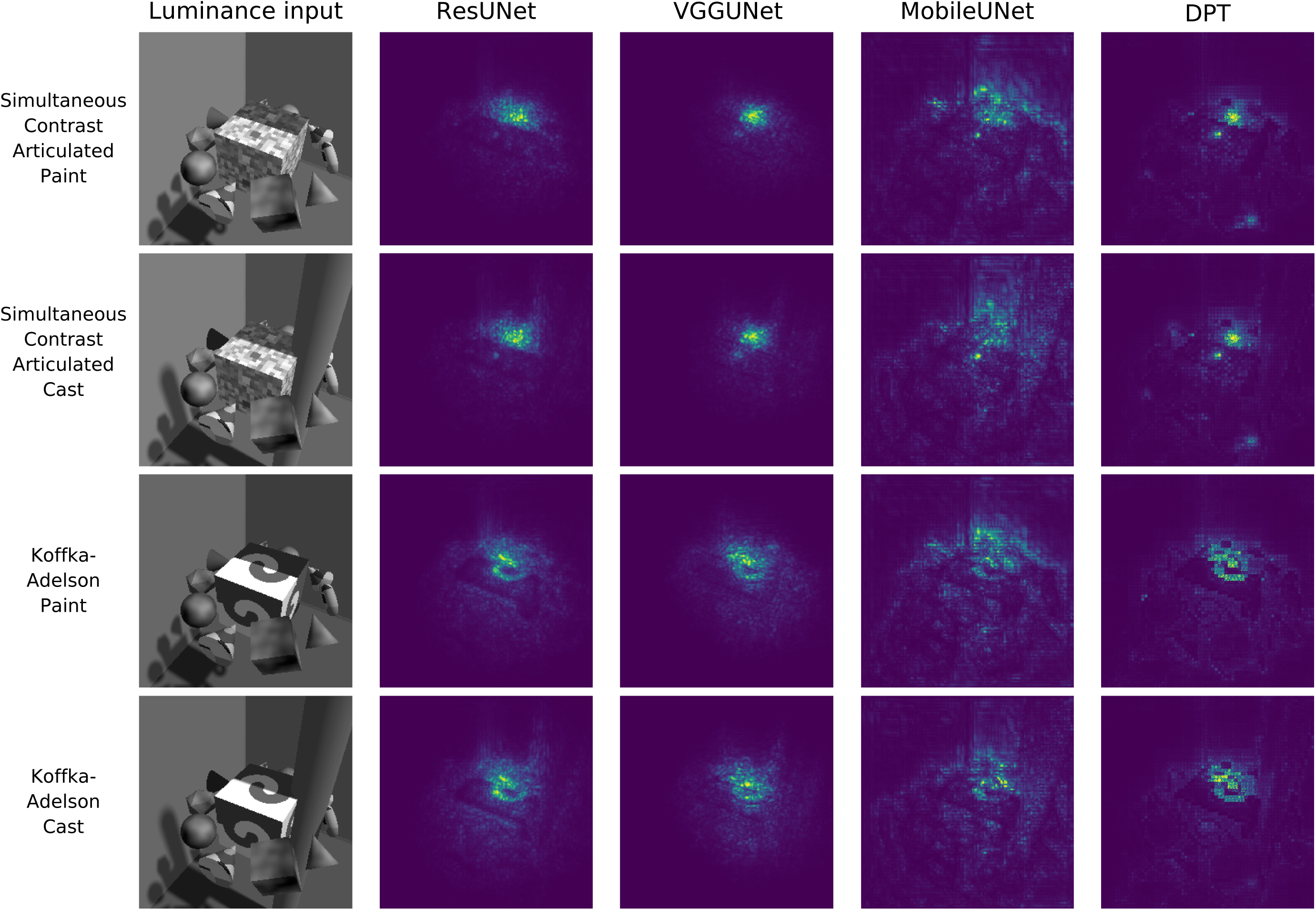
Saliency maps: illusion-based objective, frozen models.

**Figure 14.**
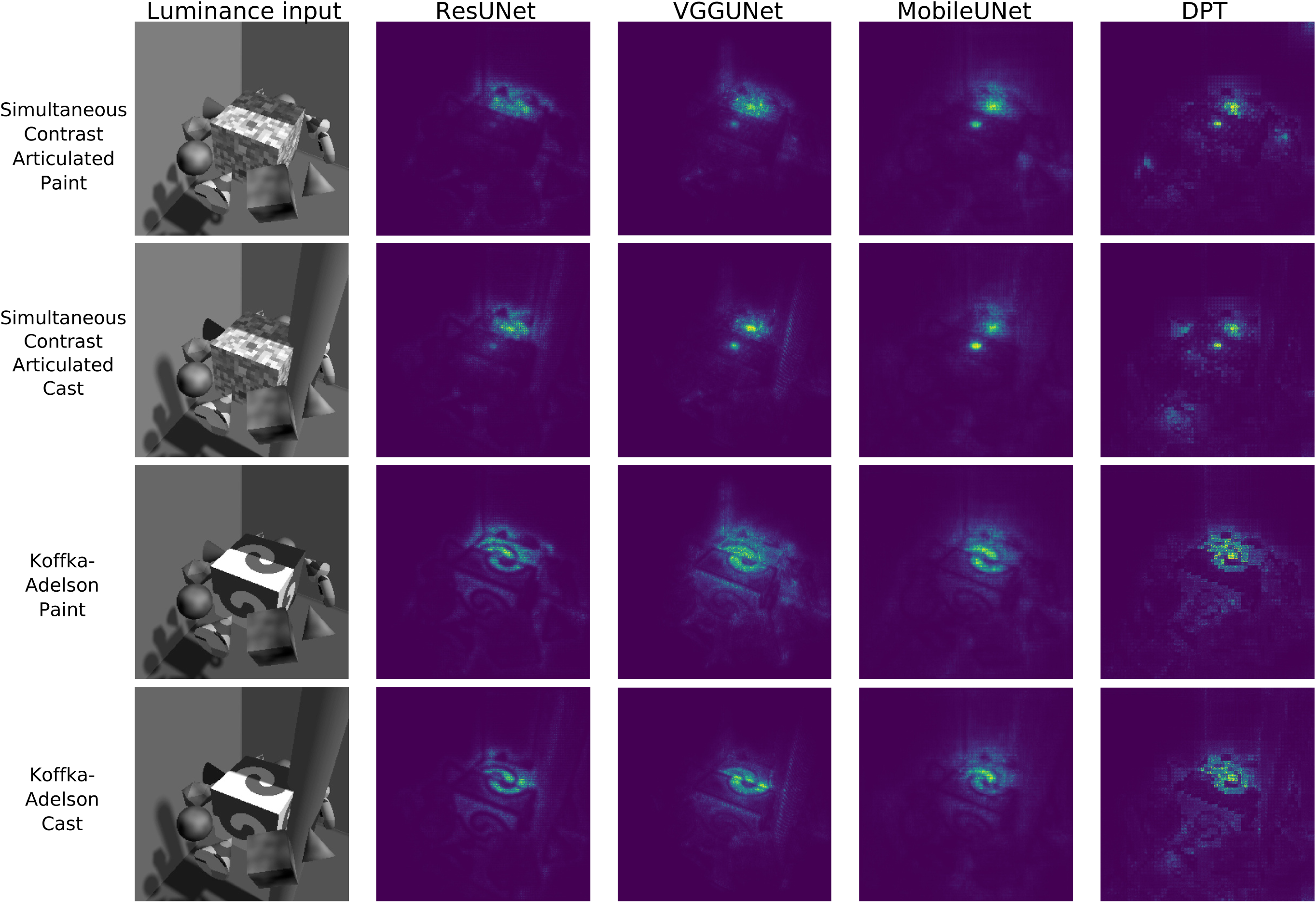
Saliency maps: illusion-based objective, fine-tuned models.

To quantify these comparisons, for each scalar objective (global and illusion-based) we computed the correlation between the saliency maps for each pair of network architectures, and averaged these correlations across illusions (Figure 15). High correlations would indicate that similar image regions are associated with reflectance estimates in the two networks. For the global objective (left panel), similarity between different networks was weak, with a mean correlation of 0.37. The lowest correlations were between CNN and DPT architectures, e.g., zero correlation for DPT vs. MobileUNet-frozen. These low correlations support the visual observation that, for the global objective, saliency maps differed substantially across architectures.

**Figure 15.**
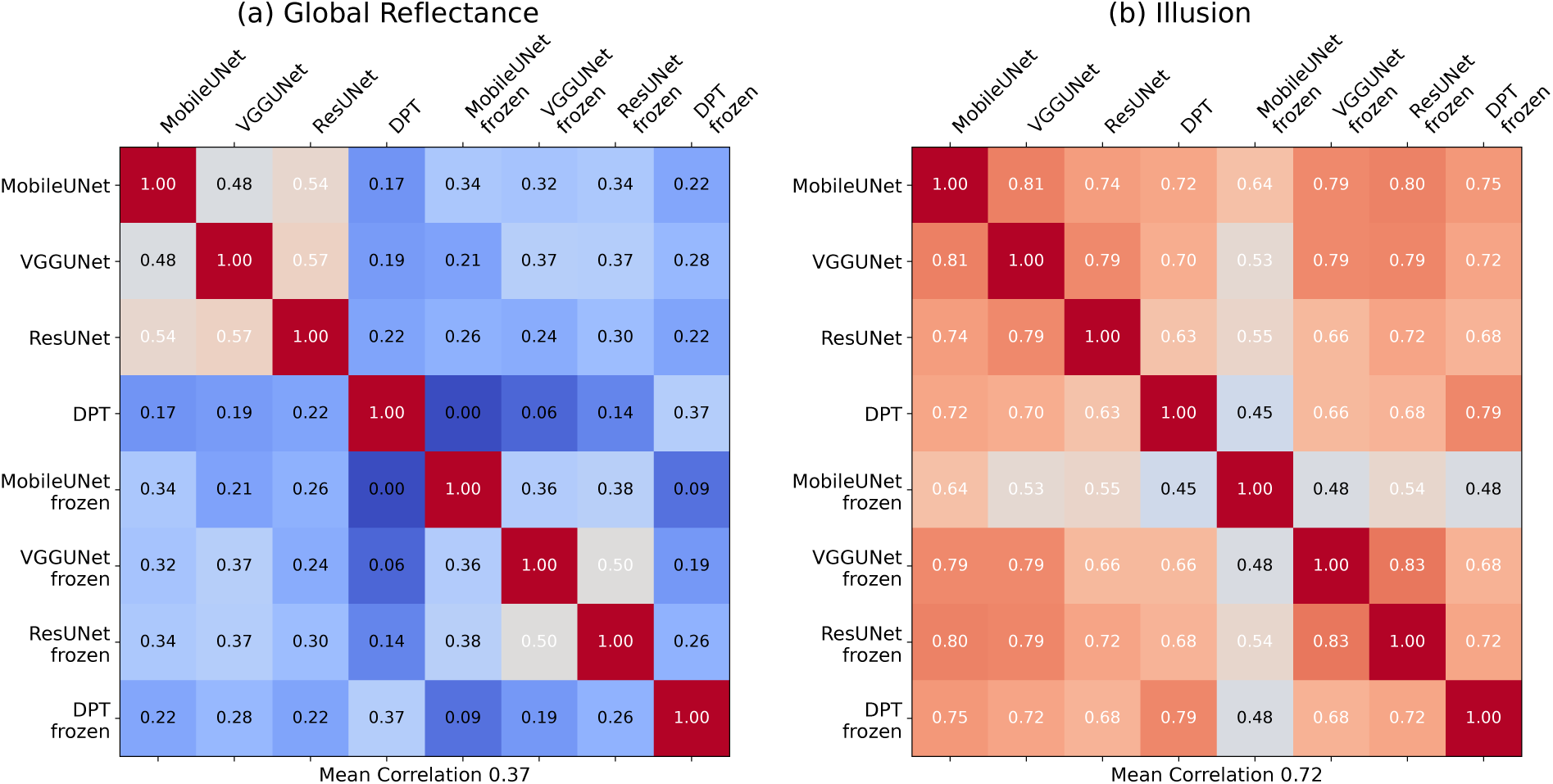
Pairwise correlations between saliency maps averaged across illusion stimuli for two scalar objectives: (a) global reflectance and (b) illusion magnitude. Rows and columns correspond to different model architectures and training conditions (frozen vs. fine-tuned encoders). The values in each cell indicate Pearson correlation coefficients. The mean correlation across model pairs is shown below each panel.

Correlations were much higher for the illusion-based objective (right panel), with a mean correlation of 0.72. As with the global objective, the lowest correlations occurred for comparisons of MobileUNet and DPT. Saliency maps for different architectures were highly correlated, as were maps for frozen and fine-tuned variants. This supports the visual impression that fine tuning and even the choice of architecture had relatively subtle effects on which image features contributed to the illusions for various networks, compared to results for the global objective.

## Discussion

Here we investigated whether deep neural networks trained for reflectance estimation mimic human lightness perception across a range of architectural choices and training constraints, and whether gradient-based saliency analyses can provide insight into differences between models. All four architectures were able to learn to estimate reflectance reasonably well, indicating that success at reflectance estimation does not depend strongly on the choice of network structure.

The networks were also susceptible to several lightness illusions, and illusion magnitudes were broadly consistent across architectures. This extends our earlier findings based on a single architecture, and shows that these results are not idiosyncrasies due to particular network features, such as convolutional processing or specific receptive field structures (Patel et al., 2026). Instead, these illusions seem to arise as a robust byproduct of inferring surface reflectance from ambiguous luminance images (Belhumeur et al., 1999).

The comparison between frozen and fine-tuned encoders provides further insight. Even models with frozen, pretrained encoders were able to learn to estimate reflectance, despite relying on features that had been optimized for completely different tasks, namely image classification and depth estimation. Furthermore, frozen-encoder models preserved not only overall performance, but also susceptibility to lightness illusions.

This shows that reflectance estimation and associated phenomena can emerge as a result of downstream specializations built on general-purpose visual features. This is consistent with the current lack of clear evidence for a dedicated reflectance-processing pathway in the human visual system, and is compatible with the idea that lightness perception may rely on general-purpose, reusable visual representations rather than early, specialized feature detectors. Fine-tuned models achieved similar or slightly improved accuracy, indicating that task-specific optimization can improve reflectance processing, even if it is not necessary.

Gradient-based saliency analyses revealed the regions where small perturbations most strongly influenced network outputs. A key finding was that the degree of similarity in saliency maps across models depended strongly on the scalar objective. Saliency maps based on mean global reflectance showed weak similarity across architectures, meaning that different architectures had very different dependencies on input regions. However, saliency maps based on illusion strength were quite similar across architectures and training protocols. The most influential stimulus regions were at and around the equiluminant regions that defined the illusions, with particular sensitivity in darker areas of the stimulus. Using classification images from human observers, Kim et al. (2018) also found that the most influential regions in a lightness judgement task were in the immediate surround of the regions being judged. Interestingly, the networks’ stronger dependence on dark image regions seems to be inconsistent with theories proposing that, in human lightness perception, the most luminous surface in a stimulus plays a key role as an ‘anchor’ for assigning reflectances (Gilchrist, 2006; Heinemann, 1955).

In the computer vision literature, there has been a useful and nuanced discussion of what can and cannot be inferred from saliency maps. Saliency maps provide a visualization of image regions that affect a model output, but they should not be understood uncritically as completely revealing the model’s decision process. Prior work has shown that some saliency methods can be insensitive to model parameters or training labels, can fail invariance tests, and can produce visually plausible maps even when they do not faithfully reflect the learned model (Adebayo et al., 2018; Hooker, Erhan, Kindermans, & Kim, 2019; Kindermans et al., 2019). The type of saliency map we used shows the magnitude of the partial derivative of a scalar function of the output, with respect to each input pixel. ‘Salient’ locations, therefore, are ones where small perturbations have large effects on the scalar function of the output. This does *not* mean that other, non-salient locations are unimportant. Completely deleting non-salient regions may have a large effect on the output, e.g., even if small perturbations to image features representing a shadow-casting object have a negligible effect on the output, those features might nevertheless play an important role in the network’s interpretation of a scene, and deleting them entirely might dramatically change the network’s output. This is a useful reminder of the fact that a saliency map, which is based on a gradient at a single point in image space, does not completely characterize how a network processes a stimulus. Even networks that have identical saliency maps may process a stimulus in fundamentally different ways, to be revealed by other methods. Thus our findings show that the architectures we examined are similar in the local perturbations that affect illusion strength, but other analyses may nevertheless reveal differences between them.

Together with our previous work (Flachot et al., 2026; Patel et al., 2026), these findings provide insight into how deep neural networks may be useful as models of human lightness perception. These models reproduce some key behavioral properties of lightness perception, such as partial constancy and susceptibility to illusions. These behaviors can emerge even from downstream decoders based on features detected by general-purpose encoders. However, the fact that we find similar results with diverse architectures suggests that the internal structure of network models provides little insight into the detailed computations underlying human lightness perception. Rather, the networks are trained to infer reflectance from ambiguous luminance cues, and the interesting result is that they then successfully reproduce some aspects of human lightness perception.

Future work could extend these analyses by comparing saliency maps with analogous methods for understanding human vision, such as classification images and gaze-contingent paradigms (Ahumada, 2002; Eckstein, 2017; Kim et al., 2018; Murray, 2011). These methods recover spatial weighting maps that show which image regions human observers rely on to make lightness judgments, and so reveal similar information to saliency maps. In addition, manipulating features of illusions embedded in three-dimensional scenes, such as local surround luminance, and illumination cues such as penumbra, would allow further characterization of what image features guide both human and model behavior (Adelson, 2000; Boyaci, Maloney, & Hersh, 2003; Radonjić, Cottaris, & Brainard, 2015). Finally, it may be possible to use physically based differentiable rendering to find stimli that are metameric for different lightness models (Beilharz & Wallis, 2026), or on which models disagree, adding an additional tool for comparing and contrasting image-computable model behavior. Together, these methods may provide further tests of neural network models of biological vision, and help to reveal computational principles underlying lightness perception.

## Author contributions

All authors contributed to the two previous papers that this work builds on (Flachot et al., 2026; Patel et al., 2026). JP and RFM developed the general plan for this paper. JP made specific choices about what networks to test, implemented all networks, ran all experiments with the networks, did all analyses of the results, and wrote the first draft of the manuscript. All authors contributed to interpreting the results and writing the manuscript.

## References

Adebayo, J., Gilmer, J., Muelly, M., Goodfellow, I., Hardt, M., & Kim, B. (2018). Sanity checks for saliency maps. In S. Bengio, H. Wallach, H. Larochelle, K. Grauman, N. Cesa-Bianchi, & R. Garnett (Eds.), Advances in neural information processing systems (Vol. 31). Curran Associates, Inc.

Adelson, E. H. (1993). Perceptual organization and the judgment of brightness. Science, 262 (5142), 2042–2044.

Adelson, E. H. (2000). Lightness perception and lightness illusions. In M. Gazzaniga (Ed.), The new cognitive neurosciences (pp. 339–351). Cambridge, MA: The MIT Press.

Ahumada, A. J. (2002). Classification image weights and internal noise level estimation. Journal of Vision, 2 (1), 121–131. doi: 10.1167/2.1.8

Allred, S. R., & Brainard, D. H. (2013). A Bayesian model of lightness perception that incorporates spatial variation in the illumination. Journal of Vision, 13(7):18, 1–18. doi: 10.1167/13.7.18

Ancona, M., Ceolini, E., Öztireli, C., & Gross, M. (2018). Towards better understanding of gradient-based attribution methods for deep neural networks. In International Conference on Learning Representations. Retrieved from https://openreview.net/pdf?id=Sy21R9JAW

Arias-Duart, A., Parés, F., Garcia-Gasulla, D., & Gimenez-Abalos, V. (2022). Focus! rating XAI methods and finding biases. In IEEE International Conference on Fuzzy Systems 2022 (pp. 1–8).

Barrow, H., Tenenbaum, J., Hanson, A., & Riseman, E. (1978). Recovering intrinsic scene characteristics. In A. R. Hanson & E. M. Riseman (Eds.), Computer vision systems (pp. 3–26). Cambridge, MA: Academic Press.

Beilharz, B., & Wallis, T. S. A. (2026). MRD: Using physically based differentiable rendering to probe vision models for 3D scene understanding. Retrieved from https://arxiv.org/abs/2512.12307v3

Belhumeur, P. N., Kriegman, D. J., & Yuille, A. L. (1999). The bas-relief ambiguity. International Journal of Computer Vision, 35 (1), 33–44.

Boss, M., Braun, R., Jampani, V., Barron, J. T., Liu, C., & Lensch, H. P. (2021). NeRD: Neural reflectance decomposition from image collections. In Proceedings of the IEEE/CVF International Conference on Computer Vision (ICCV) (p. 12684–12694).

Boyaci, H., Maloney, L. T., & Hersh, S. (2003). The effect of perceived surface orientation on perceived surface albedo in binocularly viewed scenes. Journal of Vision, 3 (8), 541–553. doi: 10.1167/3.8.2

Brainard, D. H., Longere, P., Delahunt, P. B., Freeman, W. T., Kraft, J. M., & Xiao, B. (2006). Bayesian model of human color constancy. Journal of Vision, 6(11):10, 1267–1281. doi: 10.1167/6.11.10

Brainard, D. H., & Maloney, L. T. (2011). Surface color perception and equivalent illumination models. Journal of Vision, 11(5):1. doi: 10.1167/11.5.1

Corney, D., & Lotto, R. B. (2007). What are lightness illusions and why do we see them? PLoS Computational Biology, 3 (9), e180.

Dakin, S. C., & Bex, P. J. (2003). Natural image statistics mediate brightness ‘filling in’. Proceedings of the Royal Society of London. Series B: Biological Sciences, 270 (1531), 2341–2348.

Deng, J., Dong, W., Socher, R., Li, L.-J., Li, K., & Fei-Fei, L. (2009). ImageNet: A large-scale hierarchical image database. In 2009 IEEE Conference on Computer Vision and Pattern Recognition (pp. 248–255).

DeValois, R. L., & DeValois, K. K. (1988). Spatial vision. Oxford, UK: Oxford University Press.

Eckstein, M. P. (2017). Probabilistic computations for attention, eye movements, and search. Annual Review of Vision Science, 3, 319–342.

Flachot, A., Patel, J., Wallis, T. S., Brubaker, M. A., Brainard, D. H., & Murray, R. F. (2026). Deep neural networks trained for estimating reflectance and illumination achieve lightness constancy differently than human observers. Journal of Vision, 26(20):11, 1–21. doi: 10.1167/jov.26.2.11

Gilchrist, A. (2006). Seeing black and white. New York: Oxford University Press.

Gomez-Villa, A., Martín, A., Vazquez-Corral, J., Bertalmío, M., & Malo, J. (2020). Color illusions also deceive cnns for low-level vision tasks: Analysis and implications. Vision Research, 176, 156–174.

Gomez-Villa, A., Martín, A., Vazquez-Corral, J., Bertalmío, M., & Malo, J. (2022). On the synthesis of visual illusions using deep generative models. Journal of Vision, 22(8):2, 1–18. doi: 10.1167/jov.22.8.2

He, K., Zhang, X., Ren, S., & Sun, J. (2016). Deep residual learning for image recognition. In Proceedings of the IEEE conference on computer vision and pattern recognition (pp. 770–778).

Heinemann, E. G. (1955). Simultaneous brightness induction as a function of inducing-and test-field luminances. Journal of Experimental Psychology, 50 (2), 89.

Hess, C., & Pretori, H. (1894/1970). Quantitative investigation of the lawfulness of simultaneous brightness contrast. Perceptual and Motor Skills, 31 (3), 947–969.

Hooker, S., Erhan, D., Kindermans, P.-J., & Kim, B. (2019). A benchmark for interpretability methods in deep neural networks. In H. Wallach, H. Larochelle, A. Beygelzimer, F. d’Alché-Buc, E. Fox, & R. Garnett (Eds.), Advances in neural information processing systems (Vol. 32). Curran Associates, Inc.

Howard, A. G., Zhu, M., Chen, B., Kalenichenko, D., Wang, W., Weyand, T., … Adam, H. (2017). MobileNets: Efficient convolutional neural networks for mobile vision applications. Retrieved from https://arxiv.org/abs/1704.04861

Iglovikov, V., & Shvets, A. (2018). Ternausnet: U-Net with VGG11 encoder pre-trained on ImageNet for image segmentation. Retrieved from https://arxiv.org/abs/1801.05746

Janner, M., Wu, J., Kulkarni, T. D., Yildirim, I., & Tenenbaum, J. (2017). Self-supervised intrinsic image decomposition. In I. Guyon et al. (Eds.), Advances in neural information processing systems (Vol. 30). Curran Associates, Inc.

Katz, D. (1935). The world of colour. London: Kegan & Paul.

Kim, M., Gold, J. M., & Murray, R. F. (2018). What image features guide lightness perception? Journal of Vision, 18(13):1, 1–20.

Kindermans, P.-J., Hooker, S., Adebayo, J., Alber, M., Schütt, K. T., Dähne, S., … Kim, B. (2019). The (un)reliability of saliency methods. In W. Samek, G. Montavon, A. Vedaldi, L. K. Hansen, & K.-R. Müller (Eds.), Explainable AI: interpreting, explaining and visualizing deep learning (pp. 267–280). Springer.

Kingdom, F. A. A. (2008). Perceiving light versus material. Vision Research, 48 (20), 2090–2105.

Kingdom, F. A. A. (2011). Lightness, brightness and transparency: a quarter century of new ideas, captivating demonstrations and unrelenting controversy. Vision Research, 51, 652–673.

Kingma, D. P. (2015). Adam: a method for stochastic optimization. In International Conference on Learning Representations. Retrieved from https://arxiv.org/abs/1412.6980

Koffka, K. (1935). Principles of gestalt psychology. New York: Harcourt, Brace, & World, Inc.

Krizhevsky, A., Sutskever, I., & Hinton, G. E. (2012). Imagenet classification with deep convolutional neural networks. In F. Pereira, C. Burges, L. Bottou, & K. Weinberger (Eds.), Advances in neural information processing systems (Vol. 25). Curran Associates, Inc.

Kubota, Y., Hiyama, A., & Inami, M. (2021). A machine learning model perceiving brightness optical illusions: quantitative evaluation with psychophysical data. In Proceedings of the Augmented Humans International Conference (p. 174–182). Association for Computing Machinery.

Land, E. H., & McCann, J. J. (1971). Lightness and retinex theory. Journal of the Optical Society of America, 61 (1), 1–11.

Li, Z., Shafiei, M., Ramamoorthi, R., Sunkavalli, K., & Chandraker, M. (2020). Inverse rendering for complex indoor scenes: Shape, spatially-varying lighting and svbrdf from a single image. In Proceedings of the IEEE/CVF Conference on Computer Vision and Pattern Recognition (pp. 2475–2484).

Mukherjee, A., Paul, A., & Ghosh, K. (2024). Deep learning models for perception of brightness related illusions. Applied Intelligence, 54, 10259–10283.

Murray, R. F. (2011). Classification images: A review. Journal of Vision, 11(5):2, 1–25. doi: 10.1167/11.5.2

Murray, R. F. (2020). A model of lightness perception guided by probabilistic assumptions about lighting and reflectance. Journal of Vision, 20(7):28, 1–22. doi: 10.1167/jov.20.7.28

Murray, R. F. (2021). Lightness perception in complex scenes. Annual Review of Vision Science, 7, 417–436.

Patel, J., Flachot, A., Vazquez-Corral, J., Brainard, D. H., Wallis, T. S., Brubaker, M. A., & Murray, R. F. (2026). A deep convolutional neural network trained to infer surface reflectance is deceived by mid-level lightness illusions. Journal of Vision, 26(3):14, 1–17. doi: 10.1167/jov.26.3.14

Radonjić, A., Cottaris, N. P., & Brainard, D. H. (2015). Color constancy in a naturalistic, goal-directed task. Journal of Vision, 15(13):3, 1–21. doi: 10.1167/15.3.3

Ranftl, R., Bochkovskiy, A., & Koltun, V. (2021). Vision transformers for dense prediction. In Proceedings of the IEEE/CVF international conference on computer vision (pp. 12179–12188).

Ringach, D., & Shapley, R. (2004). Reverse correlation in physiology. Cognitive Science, 28, 147–166.

Saha, S., Konkle, T., & Alvarez, G. A. (2026). A unified account of lightness illusions via edge-based reconstruction of natural images. Retrieved from https://www.biorxiv.org/content/10.64898/2026.04.07.716245v2.full.pdf

Schirillo, J. A. (2013). We infer light in space. Psychonomic Bulletin & Review, 20 (5), 905–915.

Selvaraju, R. R., Cogswell, M., Das, A., Vedantam, R., Parikh, D., & Batra, D. (2017). Grad-CAM: Visual explanations from deep networks via gradient-based localization. In Proceedings of the IEEE International Conference on Computer Vision (ICCV).

Sengupta, S., Gu, J., Kim, K., Liu, G., Jacobs, D. W., & Kautz, J. (2019). Neural inverse rendering of an indoor scene from a single image. In Proceedings of the IEEE/CVF International Conference on Computer Vision (ICCV).

Shen, L., Yeo, C., & Hua, B.-S. (2013). Intrinsic image decomposition using a sparse representation of reflectance. IEEE Transactions on Pattern Analysis and Machine Intelligence, 35 (12), 2904–2915.

Simonyan, K., Vedaldi, A., & Zisserman, A. (2014). Deep inside convolutional networks: Visualising image classification models and saliency maps. In International Conference on Learning Representations (ICLR). Retrieved from 10.48550/arXiv.1312.6034

Simonyan, K., & Zisserman, A. (2015). Very deep convolutional networks for large-scale image recognition. In International Conference on Learning Representations (ICLR). Retrieved from 10.48550/arXiv.1409.1556

Smilkov, D., Thorat, N., Kim, B., Viégas, F., & Wattenberg, M. (2017). SmoothGrad: removing noise by adding noise. Retrieved from https://arxiv.org/abs/1706.03825

Stawiaski, J. (2019). A pretrained DenseNet encoder for brain tumor segmentation. In A. Crimi, S. Bakas, H. Kuijf, B. Menze, & M. Reyes (Eds.), Brainlesion: Glioma, multiple sclerosis, stroke and traumatic brain injuries (pp. 105–115). Cham, Switzerland: Springer.

Sundararajan, M., Taly, A., & Yan, Q. (2017). Axiomatic attribution for deep networks. In Proceedings of the 34th international conference on machine learning.

Ulyanov, D., Vedaldi, A., & Lempitsky, V. (2018). Deep image prior. In Proceedings of the IEEE Conference on Computer Vision and Pattern Recognition (CVPR).

Von der Haar, L., Elvira, T., & Ochoa, O. (2023). An analysis of explainability methods for convolutional neural networks. Engineering Applications of Artificial Intelligence, 117, 105606.

White, M. (1979). A new effect of pattern on perceived lightness. Perception, 8 (4), 413–416.

Wichmann, F. A., & Geirhos, R. (2023). Are deep neural networks adequate behavioral models of human visual perception? Annual Review of Vision Science, 9, 501–524.

Yu, Y., & Smith, W. A. P. (2018). InverseRenderNet: learning single image inverse rendering. In IEEE/CVF Conference on Computer Vision and Pattern Recognition (CVPR) (pp. 3155–3164).

